# The glycan alphabet is not universal: a hypothesis

**DOI:** 10.1101/2020.01.15.908491

**Authors:** Jaya Srivastava, P. Sunthar, Petety V. Balaji

## Abstract

Several monosaccharides constitute naturally occurring glycans but it is uncertain if they constitute a universal set like the alphabets of proteins and DNA. Based on the available experimental observations, it is hypothesized herein that the glycan alphabet is not universal. Data on the presence / absence of pathways for the biosynthesis of 55 monosaccharides in 12939 completely sequenced archaeal and bacterial genomes are presented in support of this hypothesis. Pathways were identified by searching for homologs of biosynthesis pathway enzymes. Substantial variations are observed in the set of monosaccharides used by organisms belonging to the same phylum, genera and even species. Monosaccharides are grouped as Common, Less Common and Rare based on their prevalence in Archaea and Bacteria. It is observed that fewer enzymes suffice to biosynthesize the Common group. It appears that the Common group originated before the formation of three domains of life. In contrast, the Rare group are confined to a few species in a few phyla, suggesting that they evolved much later. Fold conservation, as observed in aminotransferases and SDR superfamily members involved in monosaccharide biosynthesis, suggests neo- and sub-functionalization of genes leading to the formation of Rare group monosaccharides. Non-universality of the glycan alphabet begets questions about the role of different monosaccharides in determining an organism’s fitness.

**Impact statement:** Carbohydrates, nucleic acids and proteins are important classes of biological macromolecules. The universality of DNA, RNA and protein alphabets has been established beyond doubt. However, the universality of glycan alphabet is unknown primarily because of the challenges associated with the elucidation of glycan structures. This has precluded a comprehensive investigation of glycan alphabet. To address this challenge, we have identified the prevalence of 55 monosaccharide biosynthesis pathways in 12939 completely sequenced archaeal and bacterial genomes by searching for homologs of biosynthesis pathway enzymes using HMM profiles, and in a few cases, BLASTp. This revealed that the glycan alphabet is highly variable; in fact, significant differences are found even among different strains of a species. Possible implications of this variability may be significant in understanding the evolution of Archaea and Bacteria in diverse and competitive environments. Factors that drive the choice of monosaccharides used by an organism need to be investigated, and will be of interest in understanding host-pathogen interactions. Additionally, the knowledge of glycan alphabet can be employed for structural characterization / validation of glycans inferred using mass spectrometry. Knowledge of unique monosaccharides and biosynthetic enzymes can also be used as novel drug targets against human pathogens.

**Data summary:** The curated set of proteins used in this study, with domain assignment, is listed in supplementary_data.xlsx. Corresponding 396 references with evidence of experimental characterization are included in supplementary material. Results of genome scan which include predictions of monosaccharides as well as the biosynthesis pathway enzymes is available at http://www.bio.iitb.ac.in/glycopathdb/ including the aforementioned information. Python script used to scan genomes to search for monosaccharide biosynthesis pathways are available on request.

## Introduction

Living organisms show enormous diversity in organization, size, morphology, habitat, etc., but are unified by the highly conserved processes of central dogma: replication, transcription and translation. The enormous diversity seen in life forms is encoded by DNA and decoded primarily by proteins. Both DNA and proteins use the same set of building blocks (nucleotide bases and amino acids, respectively) in all organisms; yet, they store the requisite information by merely varying the (i) set/subset of building blocks used, (ii) number of times each building block is used and (iii) sequence in which the building blocks are linked [collectively referred to as the ‘sequence’ (Table 1)]. The information required for several other biological processes are stored by glycans, the third group of biological macromolecules (1). It has been found that glycans evolve rapidly in response to changing environmental conditions, especially in Bacteria, and thus contribute to organismal diversity (2,3). The question is, do glycans use the same set of building blocks (viz., monosaccharides) in all organisms, the way proteins and nucleic acids do?

**Table 1.** Sources of diversity in primary structures of DNA, proteins and glycans.

| Feature | DNA | Protein | Glycan |
| --- | --- | --- | --- |
| *Structural diversity of building blocks* | (i) Low (four nucleotides).<br>(ii) Nucleotide modifications (known but rare): N7-methylation of Ade/Gua | (i) Higher, relative to DNA (20 amino acids). Has structurally similar pairs: Asp/Glu, Asn/Gln, Phe/Tyr, Leu/Ile/Val.<br>(ii) Amino acid modifications (known but rare): hydroxylation of Pro and Lys, selenocysteine, pyrrolysine | (i) Highest. Several pentoses and hexoses, many of which are configurational isomers.<br>(ii) Pyranose and furanose forms (e.g., Gal).<br>(iii) Both enantiomeric forms (e.g., Gal).<br>(iv) Modifications extremely common (deoxy, uronic acid, deoxyamino and its derivatives, acetylation, sulfation, ...) |
| *Linkage* | 3',5'-phosphodiester. 5',5'-phosphodiester occurs but very rare | Amide bond. γ-COOH of Glu and ε-NH<sub>2</sub> group of Lys used but very rare | Alternative isomeric linkages are very common (α1→3, β1→3, α1→6, β1→4, α2→3, α2→6 and so on) |
| *Sequence* | (i) Set/subset of building blocks used<br>(ii) Number of times each building block is used<br>(iii) Sequence in which the building blocks are linked |  |  |
| *Branching* | Absent | Absent | Quite common |
| *Sequence repeat heterogeneity* | Present | Present | Present |
| *Microheterogeneity*<sup>1</sup> | Absent | Absent | Present |

<sup>1</sup>Microheterogeneity refers to the presence of multiple forms of glycans (minor but distinct variations) present in different molecules of a protein synthesized by a cell at the ‘same’ time. This feature is unique to glycans just as the presence of splice variants is unique to proteins.

Monosaccharides show a lot more structural variation than amino acids in terms of the enantiomeric forms (both D and L), size (5 to 9 carbon atoms), ring type (pyranose, furanose), and type and extent of modification (deoxy, amino, N-formyl, N-acetyl, etc.). Some pairs of monosaccharides differ from each other merely in the configuration of carbon atoms. The sequence [as defined above] of monosaccharides brings about diversity even in the primary structure of glycans. DNA and protein are linear polymers and the linkage type that connects monomers remains the same throughout. In contrast, glycans can be branched and have alternative isomeric linkages (e.g., α1→3, β1→4, α2→6 and so on) (4), two features that enhance diversity in glycans. Repeat length heterogeneity (the number of occurrences of a sequence repeat) is observed in glycans (5,6), as well as DNA and proteins, although there are no data on the frequency of occurrence of this feature in these three classes of biomolecules. An additional factor that contributes to the diversity in the primary structure of glycans is microheterogeneity (7), a feature not seen in DNA or proteins (Table 1). These structural variations demand the use of multiple analytical techniques for sequencing and hence there are no automated methods for sequencing glycans. Biosynthesis of DNA and proteins is template-driven but not that of glycans. Consequently, there is no equivalent of polymerase chain reaction or recombinant protein expression to ‘amplify’ glycans to obtain samples in amounts required for structural / functional analysis. These constraints have largely limited data on glycan sequences.

Monosaccharides are viewed as the third alphabet of life (8). How large is this alphabet? The number of monosaccharides used collectively by living systems is at least 60. An analysis of the bacterial glycan structural data showed a distinct difference in the set of monosaccharides used by bacteria and mammals (9). Is this difference evidence of absence i.e., monosaccharides found in databases are true representations of monosaccharides used by these organisms, and those not found are not used by organisms? Or, is it just absence of evidence i.e., the glycan alphabet is indeed universal and the observed differences are merely due to inadequate sequencing? With the availability of the whole genome sequence of a large number of organisms, it has now become possible to resolve this issue.

In this study, it is hypothesized that the glycan alphabet is NOT universal, i.e., different organisms use different sets of monosaccharides. This is in contrast to those of DNA, RNA and proteins. This hypothesis is put forward based on the observations that >60 monosaccharides are found in living systems; the database of glycan structures shows differential usage of monosaccharides and that several serotypes differ from each other in the monosaccharides they use. Results obtained by mining whole genome sequences of 303 Archaea and 12636 Bacteria are presented herein in support of this hypothesis. Monosaccharides considered in this study are nucleotide activated moieties which are utilized by glycosyltransferases (GTs) in the biosynthesis of glycans. Subsequent to such a GT-catalysed transfer, monosaccharides may be modified (e.g., O-acetylation). Monosaccharide derivatives so obtained are not considered in the present study. Enzymes catalysing one or more steps of the biosynthesis pathway are not characterized experimentally for some of the monosaccharides. Such monosaccharides were not considered in this study.

## Methods

### Databases and software

Protein sequences and 3D structures were obtained from UniProt and PDB (Table S1). Completely sequenced genomes of 303 Archaea and 12636 Bacteria were obtained from the NCBI RefSeq database. These genomes are spread across 3384 species belonging to 1194 genera (Figure S1). Gene neighborhood was analyzed using feature tables taken from NCBI for the respective genomes. BLASTp, MUSCLE, HMMER and CD-Hit (Table S1) were installed and used locally. Default values were used for all parameters except when stated otherwise. Word size was set to 2 for BLASTp to prioritize global alignments over local alignments. Thresholds for Hidden Markov Model (HMM) profiles were set based on the best 1 domain bit score rather than e-values since the former is independent of database size.

### Searching genomes for monosaccharide biosynthesis pathways

Pathways for the biosynthesis of 55 monosaccharides have been elucidated to date (Table 2, Figure S2). HMM profiles were generated using carefully curated sets of homologs for 57 families of enzymes that catalyze various steps of 55 monosaccharides (Supplementary_data.xlsx:Worksheet1). Sequences were used directly as BLASTp queries when the number of enzymes characterized experimentally is not sufficient for a HMM profile (Supplementary_data.xlsx:Worksheet2). In-house python scripts were used to scan genomes to identify homologs. Presence of a homolog for each and every enzyme of the biosynthetic pathway of a monosaccharide is taken as evidence of the utilization of this monosaccharide by the organism. On the other hand, absence of a homolog for even one enzyme of the pathway is interpreted as the absence of the corresponding monosaccharide from the organism’s glycan alphabet.

**Table 2.** Summary of the pathways for the biosynthesis of monosaccharides^a,b^.

| Details about the end product of biosynthesis pathways | Precursor<sup>c</sup> |  |  |  |  |  |
| --- | --- | --- | --- | --- | --- | --- |
|  | Glc-1-P | Fru<sup>f</sup>-6-P | GDP-Man | UDP-Glc2NAc | Glc2NAc-1-P | Sed-7-P |
| Number of nucleotide sugars<sup>d</sup> | 27 | 2 | 8 | 16 | 1 | 4 |
| Number of monosaccharides<sup>e</sup> | 25 | 2 | 8 | 16 | 1 | 4 |
| Number of monosaccharides with different number of backbone carbon atoms |  |  |  |  |  |  |
| Pentose | 4 | - | - | - | - | - |
| Hexose | 21 | 2 | 8 | 13 | - | - |
| Heptulose | - | - | - | - | - | 4 |
| Nonulose | - | - | - | 3 | 1 | - |
| Number of monosaccharides of the two enantiomeric forms<sup>f</sup> |  |  |  |  |  |  |
| D | 19 | 2 | 5 | 12 | 1 | 3 |
| L | 6 | - | 3 | 4 | - | 1 |
| Number of monosaccharides of the two ring forms |  |  |  |  |  |  |
| Pyranose | 23 | 2 | 8 | 16 | 1 | 4 |
| Furanose | 2 | - | - | - | - | - |
| Number of monosaccharides with different nucleotides |  |  |  |  |  |  |
| ADP | - | - | - | - | - | 1 |
| CDP | 7 | - | - | - | - | - |
| CMP | - | - | - | 3 | 1 | - |
| GDP | - | 1 | 8 | - | - | 3 |
| TDP/dTDP<sup>g</sup> | 9 | - | - | - | - | - |
| UDP | 11 | 1 | - | 13 | - | - |

<sup>a</sup> The monosaccharide L-Iduronic acid has not been considered in this study since there is no separate pathway for its biosynthesis. Dermatan sulfate epimerase-1 or -2 (DS-epi1 or DS-epi2) catalyses C5-epimerization of glucuronic acid to L-iduronic acid in chondroitin sulfate polymeric chains (10).
<sup>b</sup> Enzymes catalysing one or more steps of the biosynthesis pathway are not characterized experimentally for some of the monosaccharides. Such monosaccharides were not considered in this study.
<sup>c</sup> Glc-6-P is the precursor for Glc-1-P (conversion catalysed by phosphoglucomutase), Fru<sup>f</sup>-6-P (catalysed by phosphoglucose isomerase) and Sed-7-P (formed in the non-oxidative phase of the pentose phosphate pathway). Fru<sup>f</sup>-6-P is the precursor of GDP-Man and UDP-Glc2NAc.
<sup>d</sup> There are two pathways for the biosynthesis of CMP-Leg5Ac7Ac, one starting from UDP-Glc2NAc and the other from Glc2NAc-1-P. Hence, the total number of nucleotide sugars will be 57 even though row sum is 58.
<sup>e</sup> L-Rhamnose and Qui4NAc are biosynthesized as both UDP- and TDP-/dTDP-derivatives. Hence, the number of monosaccharides is less than the number of nucleotide sugars by 2.
<sup>f</sup> The prefix D is omitted for D enantiomers whereas the prefix L is explicitly mentioned for L enantiomers.
<sup>g</sup> No distinction is made between TDP and dTDP in this work since literature suggests that both ribo- and deoxyribo-substrates are used by enzymes, albeit with varying extents of specificity depending upon the source organism. In fact, dTDP and TDP have been used synonymously by some authors.

### Choice of precursors

Glucose-1-phosphate, fructofuranose-6-phosphate and sedoheptulose-7-phosphate are precursors for many of the monosaccharides (Supplementary_data.xlsx:Worksheet6). Fructofuranose-6-phosphate and sedoheptulose-7-phosphate are intermediates in the glycolytic pathways viz., Embden-Meyerhof pathway and pentose phosphate pathway, respectively, and these enzymes are not considered for the search. Pathways for biosynthesis of UDP-Glc2NAc and GDP-mannose have been considered separately since Glc2NAc and mannose are glycan building blocks as well as intermediates in the biosynthesis of several other monosaccharides. Hence biosynthesis steps of UDP-Glc2NAc and GDP-mannose were excluded from those of their derivatives. An additional pathway for UDP-glucose biosynthesis was considered to analyze its ubiquity since UDP-glucose is part of both anabolic and catabolic pathways. The biosynthesis of CMP-Leg5Ac7Ac starting from N-acetyl-glucosamine-1-phosphate has also been considered because of the uncommon guanylyltransferase in the first step of the pathway.

### Generation of HMM profiles

An HMM profile was generated for each step of a biosynthesis pathway except where mentioned otherwise. Profiles were generated in two steps (Flowchart S1). The extended dataset was created to account for sequence divergence. In some cases, no additional sequences satisfying the aforementioned criteria were found, hence there is no Extend dataset. Each profile was given an annotation based on the enzyme activities of proteins that were used to generate the profile and an identifier of the format GPExxxxx; here GPE stands for Glycosylation Pathway Enzyme and xxxxx is a unique 5-digit number (Supplementary_data.xlsx:Worksheet1).

### Setting thresholds for HMM profiles

Thresholds for HMM profiles were set as described below (profile-wise details are given in Supplementary_data.xlsx:Worksheet1):

#### Using ROC curves

TrEMBL database was used to generate ROC curves. Several of the TrEMBL entries have been assigned molecular function electronically based on UniRule and SAAS (Table S1). It is assumed that these annotations are correct while generating ROC curves. True positives, false positives and false negatives were identified by comparing TrEMBL annotation with profile annotation.

#### Using bit-score scatter plots

Members of some enzyme families differ in their molecular function while retaining significant global sequence similarity e.g., C4- and C3-aminotransferases. Consequently, annotations of several TrEMBL sequences belonging to such families are incomplete e.g., DegT/DnrJ/EryC1/StrS aminotransferase family protein. In such cases, bit score scatter-plots were used to set thresholds (Figure S5). Scatter plot was also used to set threshold in case of hydrolysing and non-hydrolysing NDP-Hex2NAc C2 epimerases since many TrEMBL hits are just annotated as NDP-Hex2NAc C2 epimerases.

#### Using *T*_*exp*_ and *T*_*extend*_ as thresholds

*T*_*exp*_ or *T*_*extend*_ was used as the threshold for some profiles for one of these two reasons: (i) Sequences used to generate the profile are a subset of the sequences used to generate another profile; the latter set of enzymes has broader substrate specificity than those of the former set. For instance, sequences used for generating GPE02430 [TDP-/dTDP-4-keto-6-deoxyglucose 3-/3,5-epimerase] and GPE02530 (NDP-sugar 3-/3,5-/5-epimerase) are homologs but the former set has narrow specificity. *T*_*extend*_ was set as threshold for GPE02430 as lowering the threshold would make this profile less specific. (ii) For some profiles such as GPE50010 [nucleotide sugar formyltransferase], very few TrEMBL entries that score < *T*_*exp*_ have been assigned molecular function and hence ROC curve could not be generated.

#### The case of GPE00530

Scanning TrEMBL database with GPE00530 (Glucose-1-phosphate uridylyltransferase family 2) using the default threshold of HMMER (e-value = 10) resulted in 2693 hits with matching annotation and their scores ranged continuously from 705 to 303 bits and then from 57 to 41 bits. It was not possible to generate a ROC curve because of this discontinuity. Hence, 303 bits was set as the threshold.

### Profile annotations with broader substrate / product specificities

Many sequence homologs catalyse the “same” reaction but with (slightly) different substrate specificities. Sequence changes that confer such differential specificities are subtle and often unknown. HMM profiles of such families lack the ability to discriminate between sequences with varying substrate specificities. Two products, a major product and a minor product, are formed in certain enzyme catalysed reactions (11–13). It is possible that only the major product has been characterized while assaying an enzyme with broader substrate specificity. Another possibility is that only a subset of possible substrates has been assayed for. Hence, substrate specificities are broad in annotations of some of the profiles. As opposed to these, some profiles of aminotransferases and reductases are generated from enzymes which differ from each other with respect to the product formed viz., orientation (equatorial or axial) of the newly formed/added -OH / -NH2 group. Profile for 3,4-ketoisomerase is also of this type. UDP-GlcA decarboxylase (UXS) converts UDP-GlcA to UDP-4-keto xylose, which is further reduced to UDP-xylose. UDP-4-keto xylose is a minor product for human UXS whereas it is a major product for *E. coli* UXS (12). Both these enzymes are used to generate the profile GPE20030 (Supplementary_data.xlsx:Worksheet1).

#### Pathway steps associated with more than one HMM profile

Some steps are associated with more than one profile for one of these two reasons: (i) Non-orthologous enzymes known to catalyse the same reaction e.g., phosphomannoisomerases. (ii) Two or more profiles are generated, one with narrow and the other(s) with broad substrate specificity. Enzymes used for the former are a subset of enzymes used for the latter type of profiles e.g., aminotransferases. The process flow adopted to assign annotation for a sequence which satisfies thresholds for more than one profile is shown in Flowchart S2.

### Finding homologs using BLASTp instead of HMM profiles

HMM profiles were generated only when four or more experimentally characterized enzymes are available (two exceptions are discussed below). Global alignment and sequence similarity were used as the criteria to infer homology based on BLASTp search. The default values were set to be >= 90% query coverage and >=30% sequence similarity. However, these values were upwardly revised when query sequences belonged to homologous families that are functionally divergent (Supplementary_data.xlsx:Worksheet2). Specifically, similarity and coverage cut-offs were revised by performing an all-against-all BLASTp search of all experimentally characterized sequences of monosaccharide biosynthesis pathways.

*B. cereus* PdeG (Q81A42_1-328) is a retaining UDP-Glc2NAc 4,6-dehydratase (14). It shares higher sequence similarity with inverting UDP-Glc2NAc 4,6-dehydratases than with retaining dehydratases. The sequence of PdeG was compared with TrEMBL hits for the HMM profile of inverting UDP-Glc2NAc 4,6-dehydratases (GPE05331), based on which the sequence similarity cut-off for PdeG was set to 70%. The threshold for GPE05331 was set such as to exclude PdeG (Figure S5).

#### Criteria for finding homologs of UDP-2,4-diacetamido-2,4,6-trideoxy-β-L-altrose hydrolase and UDP-4-amino-6-deoxy-Glc2NAc acetyltransferase

Four experimentally characterized enzymes are known for each of these two families. However, BLASTp approach was used instead of generating an HMM profile. This is because a suitable bit score threshold could not be arrived, which in turn, was because several of the TrEMBL entries obtained as hits are annotated as CMP-N-acetylneuraminic acid synthetase or equivalent (for hydrolase), or O-acetyltransferase or equivalent (for acetyltransferase).

### Uncertainties in prediction

Any description of molecular function of a protein is stratified and includes specifying the type of reaction catalysed, substrate(s) used, etc. A vast majority of sequences conceptually translated from genome sequences are assigned molecular function based on sequence homology to experimentally characterized proteins. Even though experimental validation is available for only a small fraction of proteins due to practical constraints, such studies have shown that homology-based assignments are generally valid and deviations typically pertain to the extent of substrate specificity, metal ion dependency and such. Nevertheless, caution is warranted with increasing sequence divergence and one has to be on the lookout for homologs that have acquired new molecular function as a result of mutation of a handful of key residues (neo-functionalization). In view of this, in the present study, HMM and BLASTp thresholds have been chosen with higher stringency and assignment of substrate(s) and product(s) has been made conservatively by manually curating false positives and false negatives from the Swiss-Prot database, details of which are given below:

1. Both GDP-rhamnose and GDP-6-deoxytalose are assigned as products of the same pathway, because their biosynthesis proceeds through the same pathway with the exception of the last step being catalysed by homologous 4-reductases. It is not possible to infer if product specificity of enzymes in this family is absolute or partial i.e., one is a major product and other, a minor product, due to inadequate experimental data. An identical situation is seen in the pathways for the biosynthesis of CDP-cillose and CDP-cereose, and for CDP-abequose and CDP-paratose. In view of this, prevalence data will be the same for the two monosaccharides of a pair (Supplementary_data.xlsx:Worksheet3).
2. Non-hydrolyzing NDP-Hex2NAc C2-epimerases (GPE02030) are part of biosynthesis pathways of different monosaccharides. The extent of substrate specificity of the experimentally characterized members of this family is not known since not all enzymes have been assayed using all possible substrates. In literature, substrate specificity is arrived at based on the genomic context and the same approach has been followed in the present study as well. For example, hits for GPE02030 profile are treated as Man2NAc synthesis pathway enzymes, unless other enzymes of L-Fuc2NAc, L-Qui2NAc or Man2NAc3NAcA pathway are also present.
3. Some monosaccharides are precursors for other monosaccharides and hence, genomes predicted to have the pathway for the latter monosaccharide will also have the precursor monosaccharide. Following are the precursor-final product monosaccharide pairs encountered in this study: (i) L-Rha2NAc → L-Qui2NAc, (ii) L-Rhamnose → 6-Deoxy-L-talose, (iii) Fucose → Fucofuranose, (iv) Paratose → Tyvelose, (v) Galactose → Galactofuranose, (vi) GlcA → GalA, (vii) L-Ara4N → L-Ara4NFo, (viii) Per → Per4Ac, (ix) Man2NAc → Man2NAcA, (x) Glc2NAcA → Gal2NAcA, and (xi) Bac2Ac4Ac → Leg5Ac7Ac.
4. The pathway for the synthesis of L-arabinose is an extension of the pathway for the synthesis of xylose. However, most genomes predicted to have xylose pathway also have L-arabinose pathway. This is because UDP-sugar C4-epimerase family members (GPE02230) catalyse C4-epimerization of glucose, GlcA, Glc2NAc, Glc2NacA and xylose. Assigning substrate specificity solely based on sequence similarity is not possible. The challenge is compounded by the fact that some of these enzymes show broad substrate specificity while the rest are only specific to a single substrate. Not all enzymes have been assayed for all potential substrates.

## Results

### Glycan alphabet size is not the same across Archaea and across Bacteria

The number of monosaccharides used by different species is significantly different (Figure 1) and is independent of proteome size (Figure S3). Data for the prevalence of monosaccharides in 12939 genomes is very similar to that in 3384 species (Figure S4) indicating that the outcome is not biased by the skew in the number of genomes (strains) sequenced for a given species (Figure S1). In fact, none of the organisms use all 55 monosaccharides: the highest number of monosaccharides used by an organism is 23 [*Escherichia coli* 14EC033]. Just 1 and 2 monosaccharides are used by 188 and 117 species, respectively. Glucose, galactose and mannose, and their 2-N-acetyl (Glc2NAc, Gal2NAc, Man2NAc) and uronic acid (GlcA, GalA, Glc2NAcA, Gal2NAcA) derivatives are the most prevalent besides L-rhamnose, as the biosynthesis pathways for these monosaccharides are found in >50% of genomes (Figure 2). These monosaccharides are thus categorized as ‘Common’ group. However, none of them are used by all organisms (Supplementary_data.xlsx:Worksheet3).

**Figure 1.**
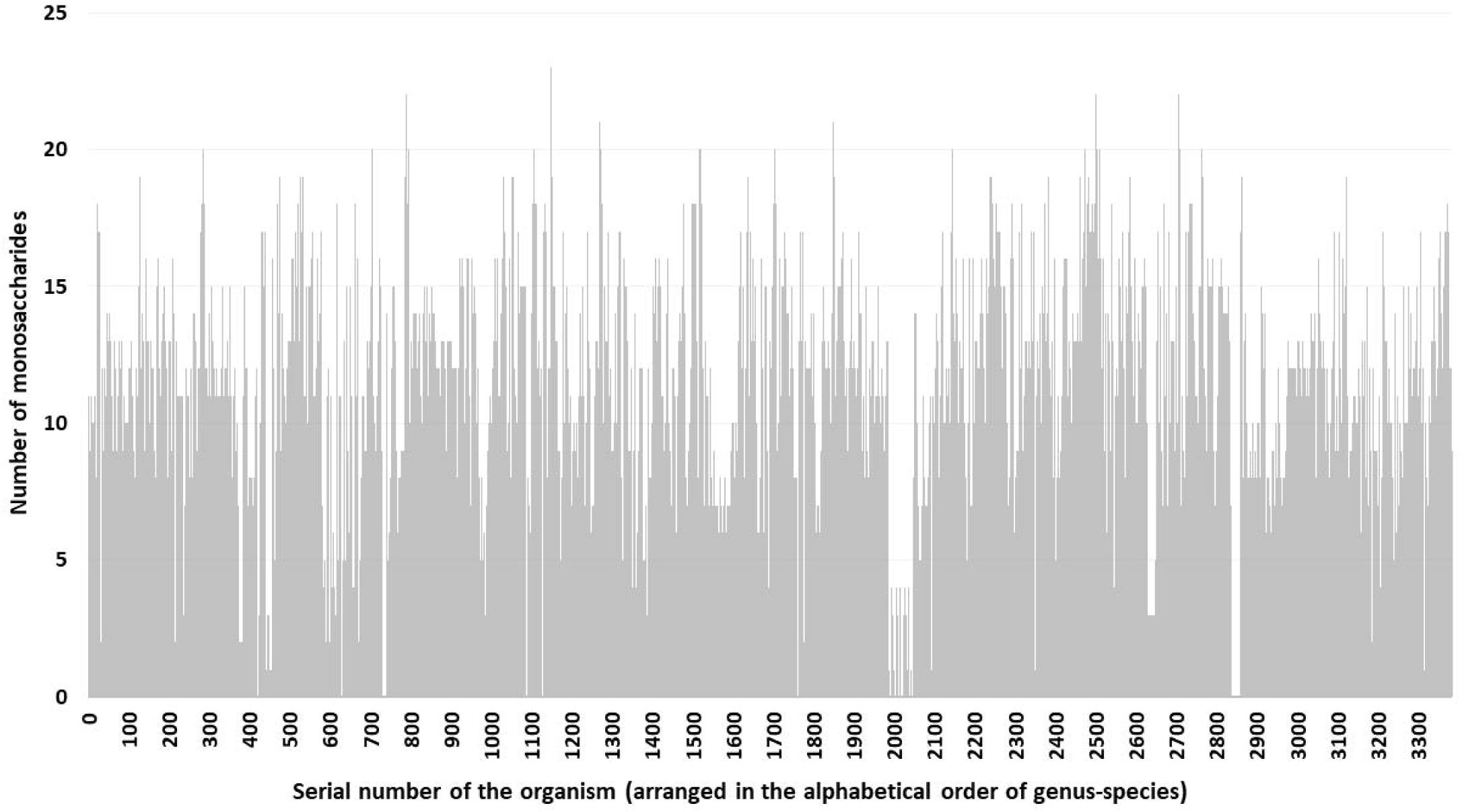
The number of monosaccharides for which biosynthesis pathways are found in a species. More than one strain is sequenced for several species (Figure S1). In such cases, data for the strain which has the highest number of monosaccharides is plotted. Total number of species = 3384.

**Figure 2.**
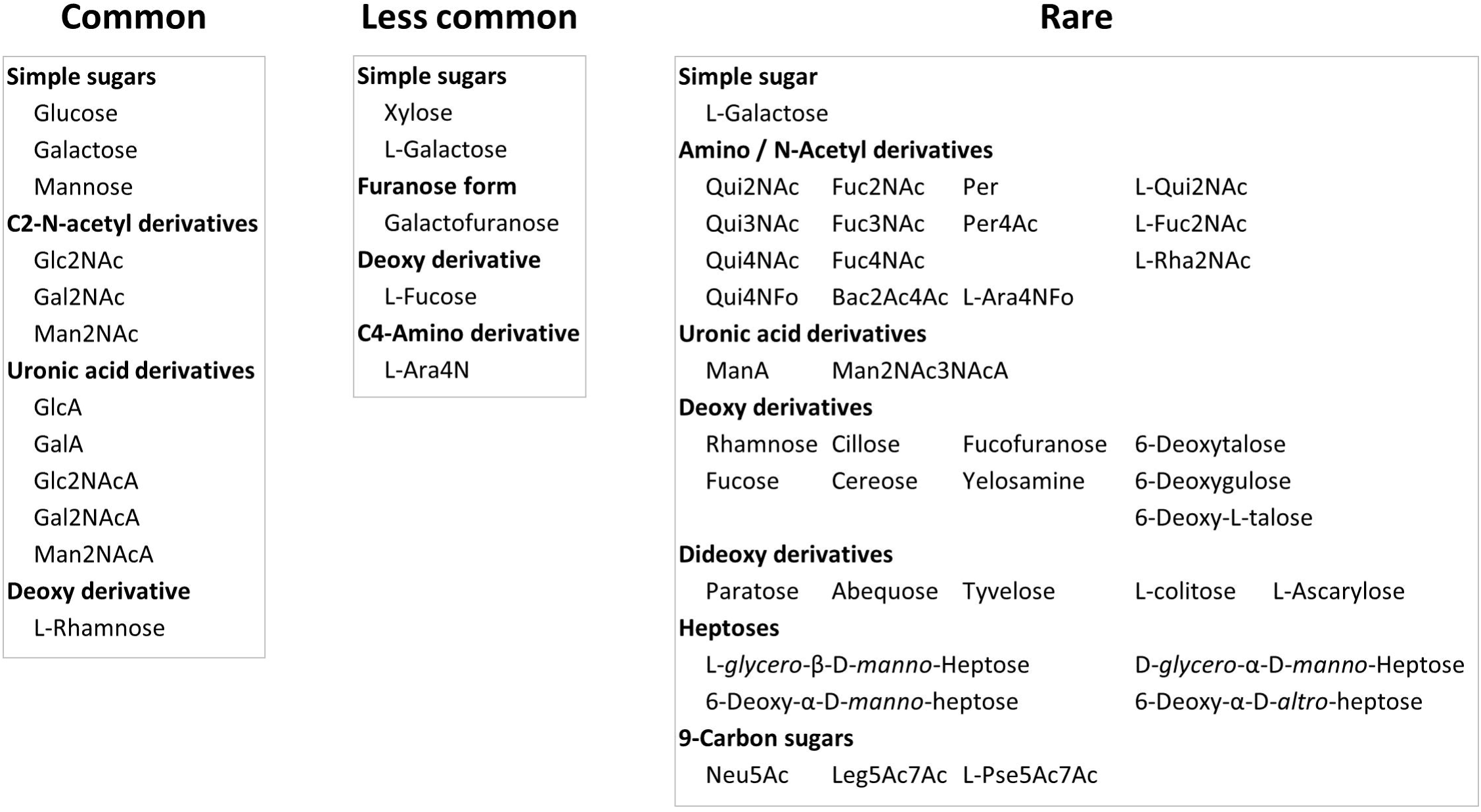
Classification of monosaccharides into three groups based on prevalence in archaeal+bacterial genomes. These groups are Common (found in >=50% of genomes), Less Common and Rare (found in <=10% of genomes). Abbreviated names are used for some of the monosaccharides. Full names of these are given in Supplementary_data.xlsx:Worksheet4.

### Evolution and diversification of glycan alphabet

It is observed that only a limited set of enzymes suffice to biosynthesize the Common group monosaccharides e.g., nucleotidyltransferases (activation), amidotransferase and N-acetyltransferase (Hex2NAc from a hexose), C4-epimerase (Glc to Gal) and C6-dehydrogenase (uronic acid) belonging to the SDR superfamily, non-hydrolyzing C2-epimerase (Glc2NAc to Man2NAc), mutase (6-P to 1-P) and isomerase (pyranose to furanose) (Figure 3 and Supplementary_data.xlsx:Worksheet5). Using this limited set of monosaccharides, organisms seem to achieve structural diversity by mechanisms such as alternative isomeric linkages, branching and repeat length heterogeneity. Some organisms use an additional set of monosaccharides, viz., L-fucose, galactofuranose, xylose, L-Ara4N and L-arabinose. These monosaccharides are categorized as Less Common group. Organisms using this group of monosaccharides have enhanced the glycan repertoire by acquiring C3/C5-epimerase, 4,6-dehydratase, C4-reductase, C6-decarboxylase and C4-aminotransferase. The rest of the monosaccharides are used by very few organisms and thus constitute the ‘Rare’ group (Figure 2).

**Figure 3.**
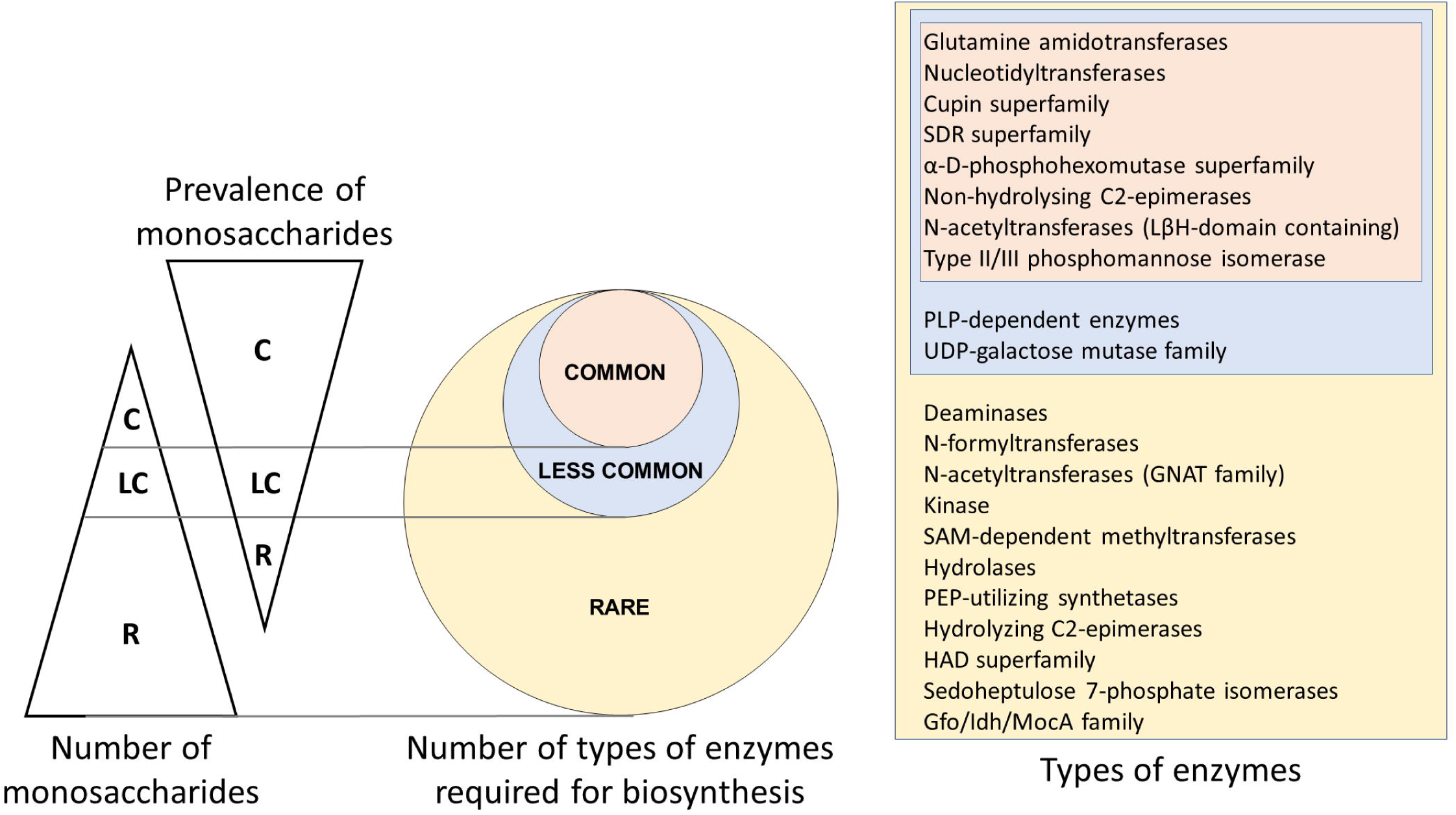
A qualitative comparison of the number of monosaccharides of the three groups viz., Common (C), Less Common (LC) and Rare (R) with their prevalence in archaeal+bacterial genomes and the number of types of enzymes required for their biosynthesis. The size of a group is inversely related to the prevalence of the corresponding group of monosaccharides. Enzymes required for the biosynthesis of Common group monosaccharides are required for the biosynthesis of Less Common and Rare groups also; similarity, those for the Less Common group are required for the biosynthesis of Rare group also. Different enzymes belonging to each of the superfamily mentioned above are listed in the file Supplementary_data.xlsx:Worksheet5. Note that the group sizes are not to scale. It may be noted that additional types of enzymes may have to be included when experimental data about the pathways for the biosynthesis of other monosaccharides becomes available. HAD, haloalkanoic acid dehalogenase Gfo/Idh/MocA, glucose_fructose oxidoreductase/inositol 2_dehydrogenase/rhizopine catabolism protein MocA GNAT, GCN5-related N-acetyltransferases LβH, left handed β helix PEP, phosphoenolpyruvate PLP, pyridoxal 5’-phosphate SAM, S-adenosyl-L-methionine SDR, short chain dehydrogenase reductase UDP, uridine diphosphate

Occurrence of the Common group of monosaccharides in all three domains of life points to their presence early on during evolution. Neo- and sub-functionalization of horizontally acquired and duplicated genes during the course of evolution have been widely reported (e.g.,(15,16)). It is envisaged that the enzymes required for the biosynthesis of Rare group monosaccharides have arisen by such neo- and sub-functionalization. Aminotransferase and short-chain dehydrogenase reductase (SDR) superfamily enzymes involved in the biosynthesis of monosaccharides lend support to this inference. Superposition of a few C3- and C4-aminotransferases show remarkable conservation of the 3D structures despite differences in the pyranose ring position at which the amino group is transferred as well as the nucleotide sugar substrate (Figure 4). 3D structures are conserved even among SDR superfamily enzymes despite catalysing different reactions viz., epimerization (at C2 or C4), removal of water (dehydratase at C4, C6) and reduction (at C4).

**Figure 4.**
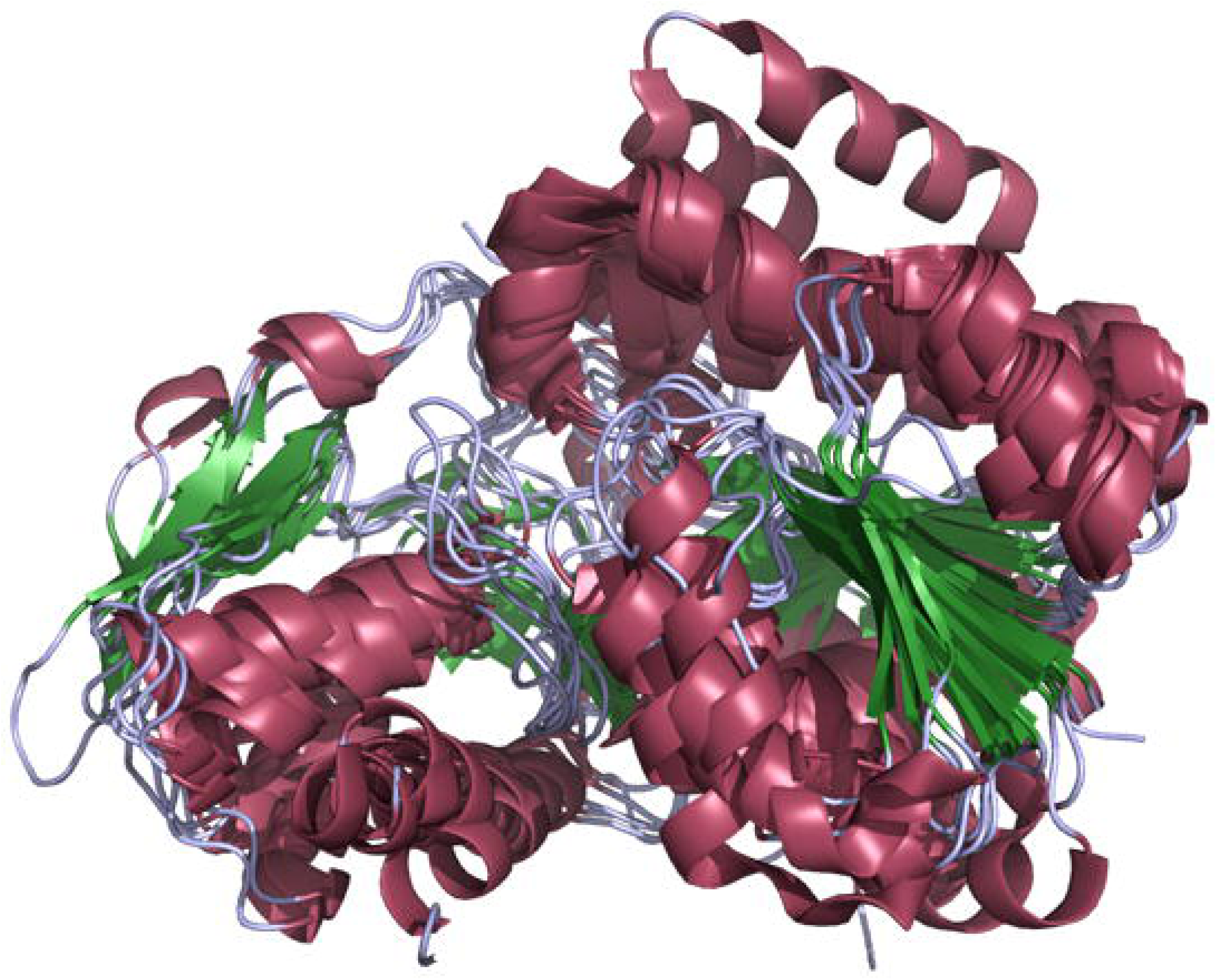
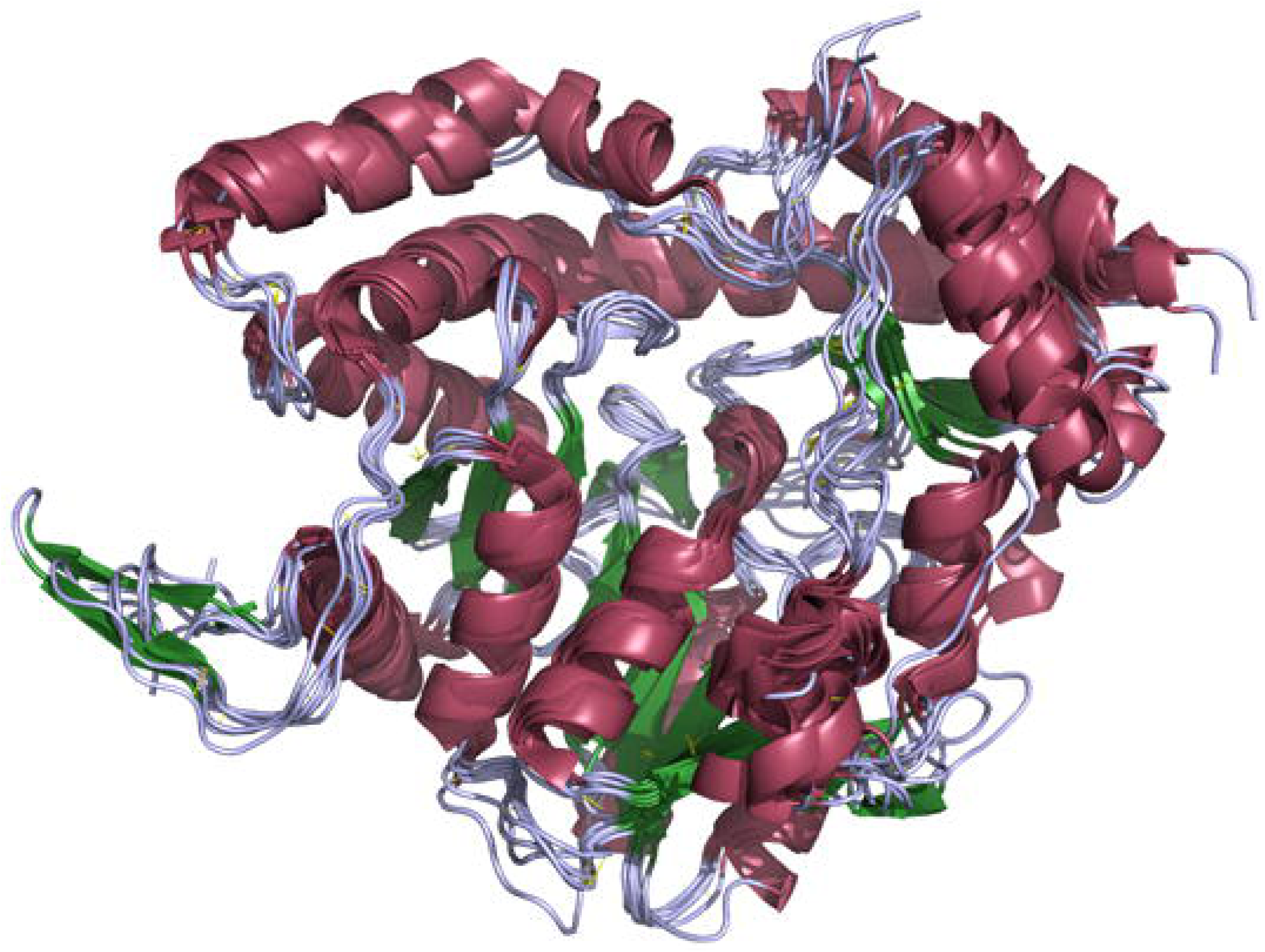
3D structural superimposition of enzymes belonging to aminotransferase (A) and SDR (B) superfamilies involved in the biosynthesis of monosaccharides. Color scheme: helices, raspberry red; sheet, forest green; loops, light blue. **Panel A**: Aminotransferase superfamily enzymes: 1MDO_A: ArnB from UDP-L-Ara4N biosynthesis; 2FNI_A: PseC from CMP-L-Pse45Ac7Ac biosynthesis; 2OGA_A: DesV from TDP-/dTDP-desosamine biosynthesis; 3BN1_A: perA from GDP-per biosynthesis; 3NYU_A: WbpE from UDP-Man2NAc3NAcA biosynthesis; 4PIW_A: WecE from TDP-/dTDP-Fuc4NAc biosynthesis; 4ZTC_A: PglE from CMP-Leg5Ac7Ac biosynthesis; 5U1Z_A: wlaRG from TDP-/dTDP-Fuc3NAc/Qui3NAc biosynthesis. ArnB, PseC, perA, WecE and PglE are C4-aminotransferases whereas DesV, WbpE and wlaRG are C3-aminotransferases. **Panel B**: 1ORR_A: RfbE, C2-epimerase from CDP-tyvelose biosynthesis; 2PK3_A: Rmd, 4-reductase from GDP-rhamnose biosynthesis; 1KBZ_A: rmlD, C4-reductase from TDP-/dTDP-L-rhamnose biosynthesis; 1T2A_A: gmd, C4,C6-dehydratase from GDP-L-fucose biosynthesis; 1SB8_A: WbpP, C4-epimerase from UDP-Gal2NAc biosynthesis; 5BJU_A: PglF, C4,C6-dehydratase from UDP-Bac2Ac4Ac biosynthesis.

### Glycan alphabet varies even across strains

Remarkably, variations in the size of glycan alphabet are significant even at the strain level (Figure 5). Strain-specific differences are pronounced in species such as *E. coli, Pseudomonas aeruginosa* and *Campylobacter jejuni* (Figure 6) possibly reflecting the diverse environments that these organisms inhabit. Among organisms which inhabit the same environment, strain-specific differences show mixed pattern: among the 71 strains of *Streptococcus pneumoniae*, the maximum and minimum number of monosaccharides utilized by a strain are 4 and 12, respectively. Such a variation could have evolved as a mechanism to evade host immune response. In contrast, strains of *Streptococcus pyogenes* and strains of *Staphylococcus aureus* inhabit the same environment (respiratory tract and skin, respectively) and show very little variation in the monosaccharides they use. Both are capsule producing opportunistic pathogens suggesting that they might bring about antigenic variation by variations in linkage types, branching etc. (17), even with the same set of monosaccharides. Strains of *Mycobacterium tuberculosis, Brucella melitensis, Brucella abortus* or *Neisseria gonorrhoeae*, all of which are human intracellular pathogens, also show insignificant variation. It is possible that different strains of a pathogen are a part of distinct microbiomes and microbial interactions within the biome/with the host determine the glycan alphabet of the organism. Availability of additional characteristics such as phenotypic data and temporal variations in glycan structures is critical for understanding the presence/absence of strain-specific variations.

**Figure 5.**
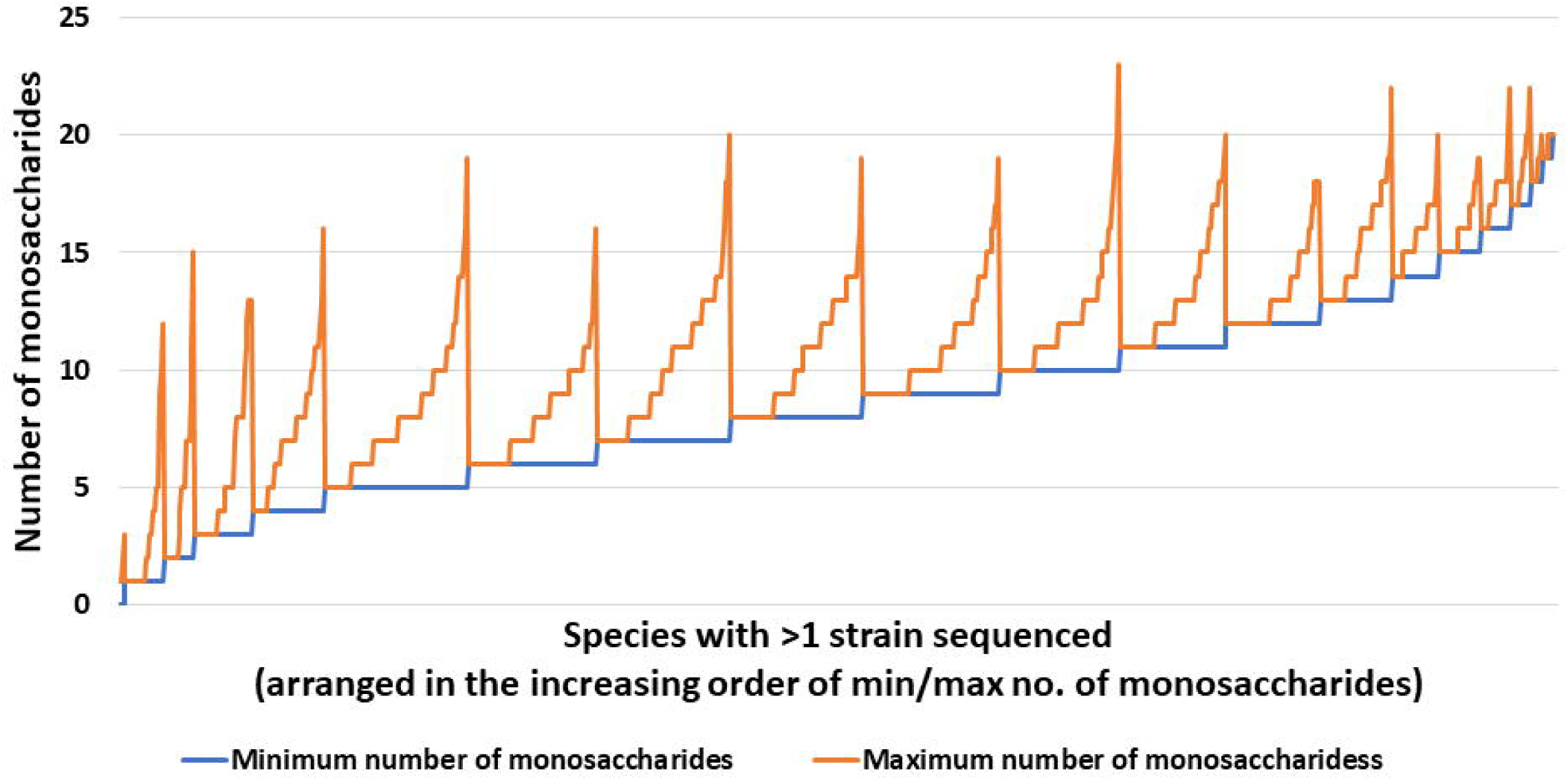
Variations in the number of monosaccharides used by different strains of a species. Species with more than one sequenced strain and at least one monosaccharide predicted in one of the strains are considered. Only the smallest and largest numbers are shown.

**Figure 6.**
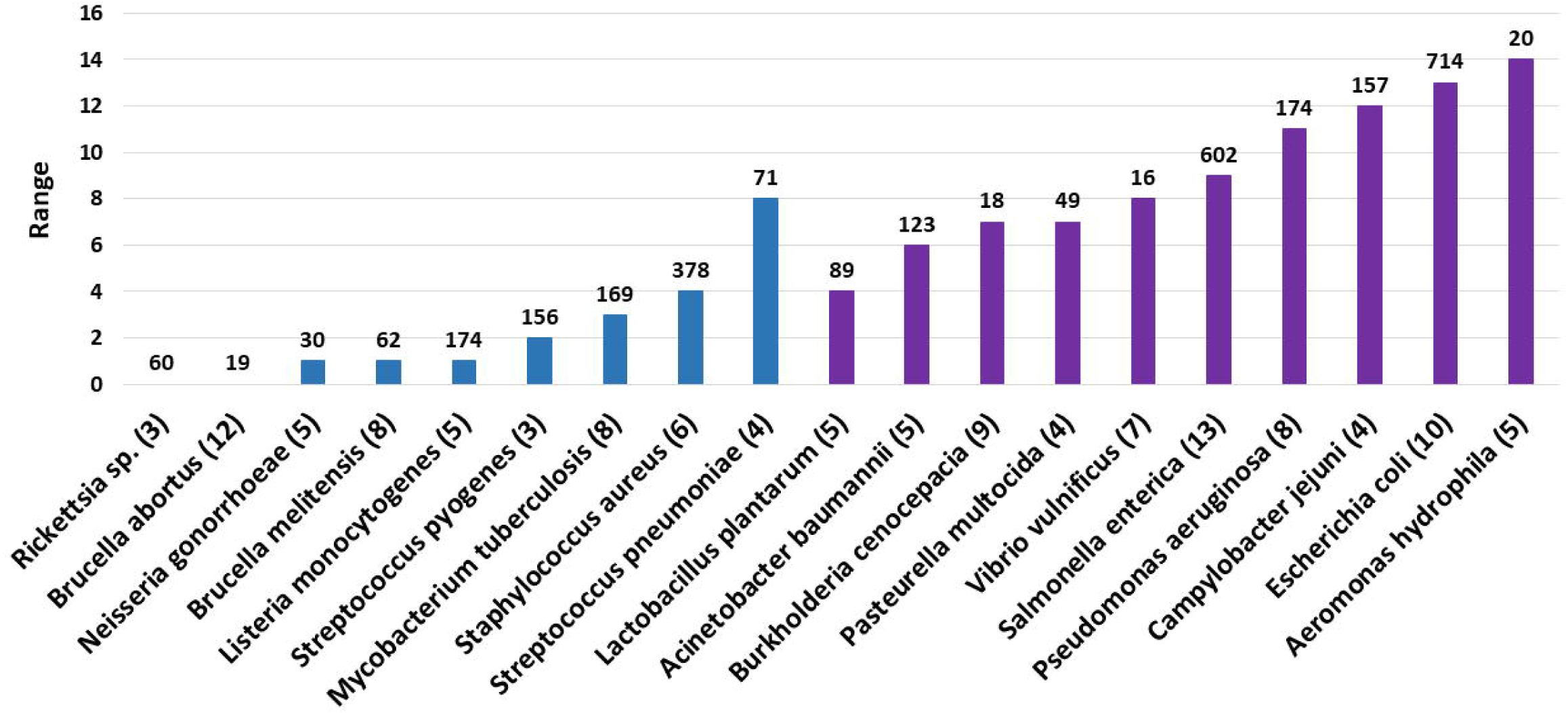
Different strains of some of the species do not use the same number of monosaccharides. The range of the number of monosaccharides used by various strains of some of the clinically important species are shown here. The number of sequenced strains for each organism is shown above the corresponding bar. Number in parenthesis after the name of each organism represents the minimum number of monosaccharides used by one of the strains of this organism. Note that the set of monosaccharides encoded by different strains utilizing the same number of monosaccharides may vary. Organisms associated with narrow habitat are shown in blue, while those with broad habitat are shown in purple.

### Prevalence of monosaccharides across Phyla

Not all sugars of the Common group (Figure 1) are found across all phyla whereas Neu5Ac belonging to the Rare group is found across all phyla. GlcA and GalA (Common group) are absent in Thermotogae suggesting that pathways for their biosynthesis are lost in this phylum. A similar conclusion is drawn for the absence of L-fucose and L-colitose in TACK group phylum. Most of the Rare group sugars are limited to very few species in a few phyla (Figure 7). For instance, Fuc4NAc and L-*glycero*-β-D-*manno*-heptose (ADP-linked) are found only in Gamma-proteobacteria, a class that comprises of several pathogens. The other three heptoses, which are GDP-linked, are absent in Gamma-proteobacteria. Recently, it was found that *Helicobacter pylori*, belonging to the class Epsilon-proteobacteria, synthesizes ADP-*glycero*-β-D-*manno*-heptose for activating the NF-кβ pathway in human epithelial cells (18). This pathway has been experimentally characterized in very few organisms. Consequently, homologs for this pathway are found by BLASTp queries and not by HMM profiles. In the present study, this pathway turned out to be a false negative because of the high stringency set for BLASTp thresholds. In view of this, it is possible that such sugars which appear restricted to a few phyla are also found in others.

**Figure 7.**
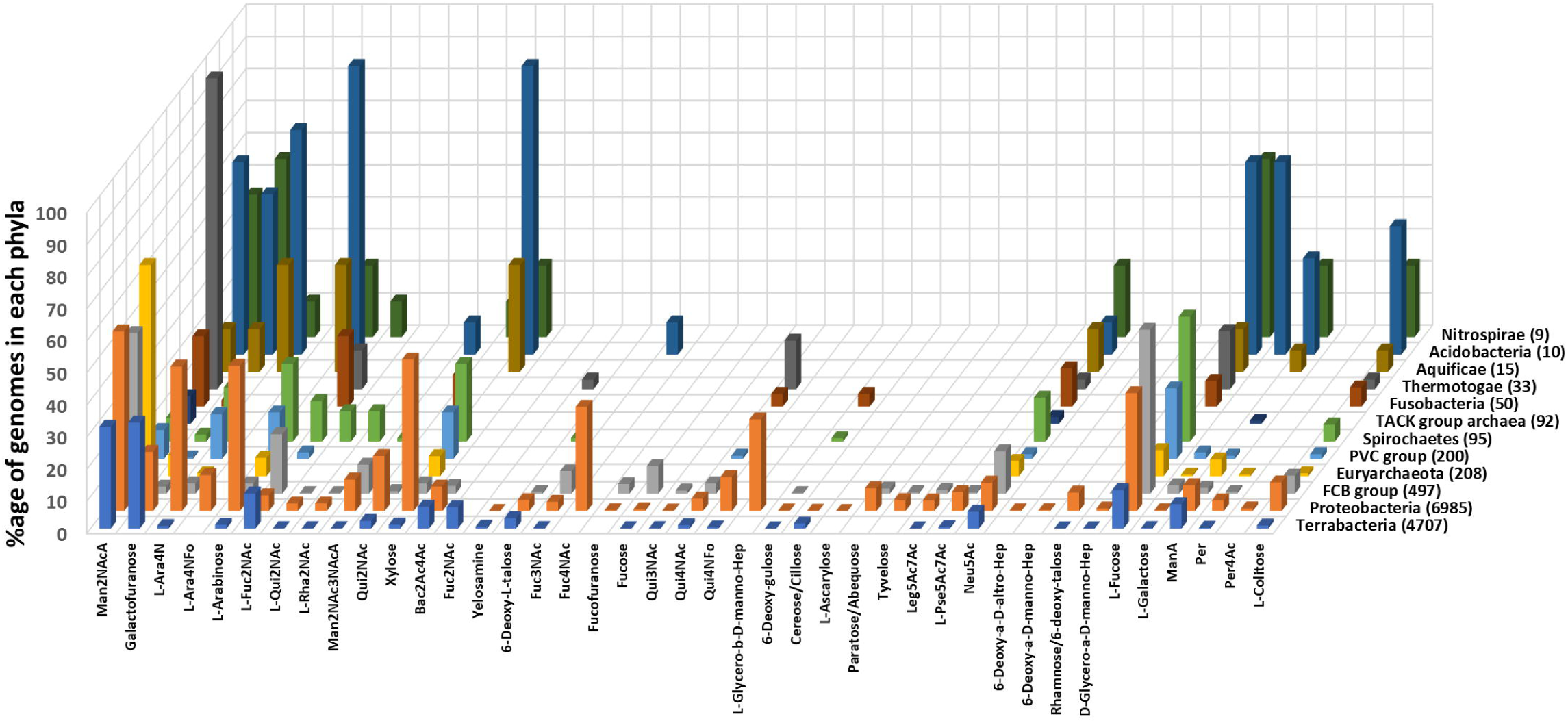
Prevalence of Less Common and Rare group monosaccharides in different microbial phyla. Data for phyla with less than five sequenced genomes are not shown to avoid visual clutter. Only names of monosaccharides are used for annotation even though all are biosynthesized as nucleotide sugars. Abbreviated names are used for some of the monosaccharides. Full names of these are given in Supplementary_data.xlsx:Worksheet4.

### Why some eubacteria do not biosynthesize any monosaccharide?

None of the monosaccharides are biosynthesized by some mollicutes (e.g., Mycoplasma) and endosymbionts (e.g., *Ehlrichia sp.* and *Orientia sp.*) because the biosynthesis pathways are completely absent. Mollicutes lack cell wall (19) which could explain the absence of monosaccharides. Endosymbionts have reduced genomes which is seen as an adaptation to host dependence (20) (21). Biosynthesis pathway enzymes are lost / are being lost as part of the phenomenon of genome reduction. This is illustrated by the endosymbiont *Buchnera aphidicola*: 13 of the 25 strains have the pathway for the biosynthesis of UDP-Glc2NAc, 7 have partial pathway and 5 do not encode any gene of this pathway. Pathway for none of the other monosaccharides are found in this organism. Pathways are incomplete i.e., enzymes catalysing one or more steps of the pathway are absent in some organisms. Some species of Mycoplasma, Ureaplasma and Spiroplasma lack mannose-1-phosphate guanylyltransferase because of which GDP-mannose is not biosynthesized. GlmU which converts Glc2N-1-phosphate to UDP-Glc2NAc is absent in *Chlamydia sp.* However, Glc2N is found in the LPS of *Chlamydia trachomatis* (22). Whether this is indicative of the presence of a transferase which uses Glc2N-1-phosphate instead of UDP-Glc2N needs to be explored.

### Do *Rickettsia sp.* and *Chlamydia sp.* source monosaccharides from their host?

*Rickettsia sp.* (60 strains), *Orientia tsutsugamushi* (7 strains), and *Chlamydia sp.* (143 strains) are obligate intracellular bacteria. *O. tsutsugamushi* does not contain pathways for the biosynthesis of any of the monosaccharides. This is in consonance with the finding that it does not contain extracellular polysaccharides (21). Rickettsia species have pathways for the biosynthesis of Man2NAc, L-Qui2NAc and L-Rha2NAc. L-Rha2NAc is the immediate precursor for L-Qui2NAc (Figure S2e). Rickettsia are known to use Man2NAc and L-Qui2NAc but not L-Rha2NAc (23) implying that UDP-L-Rha2NAc is just an intermediate in these organisms. The pathway for the biosynthesis of UDP-Glc2NAc, precursor for these Hex2NAcs, is absent suggesting partial dependence on host (human). Notably, genes for the biosynthesis of Man2NAc and L-Qui2NAc have so far not been reported in humans, which explains why Rickettsia have retained these pathways (the human genome was scanned and these pathways are not found; unpublished data). Both Rickettsia and Orientia belong to the same order, Rickettsiales. Symptoms caused by these two are similar (24). In spite of similarities in host preference and pathogenicity, *Rickettsia sp.* continues to use certain monosaccharides while diverging from *O. tsutsugamushi* (25) which uses none. Is this because Rickettsia use ticks as vectors whereas Orientia use mites (26)? *Rickettsia akari*, the only Rickettsial species which uses mites as vectors and contains pathways for Man2NAc and L-Qui2NAc biosynthesis, has been proposed to be placed as a separate group because its genotypic and phenotypic characteristics are intermediate to those of Orientia and Rickettsia (26).

### Absence of Glc2NAc in organisms other than endosymbionts

UDP-Glc2NAc is the precursor for the biosynthesis of several monosaccharides (Figures S2e, S2f). However, pathways for its biosynthesis are absent in ∽10% of the genomes excluding endosymbionts. None of the organisms in FCB group and Spirochaetes contain this monosaccharide. Further analysis revealed the loss of first (GlmS) or last (GlmU) enzyme of the pathway in several of their genomes. This pattern suggests that organisms of this phyla are in the process of losing UDP-Glc2NAc pathway. Incidentally, some of these genomes do contain its derivatives. They include host-associated organisms such as *Bacteriodes fragilis, Flavobacterium sp., Tannerella forsythia, Akkermansia muciniphila, Bifidobacterium bifidum, Leptospira interrogans*, etc., suggesting that they obtain Glc2NAc from their microenvironment. However, a few free-living organisms which contain derivatives of UDP-Glc2NAc but not UDP-Glc2NAc were also identified. For instance, GlmU is not present in *Arcticibacterium luteifluviistationis* (arctic surface seawater) and its C-terminus (acetyltransferase domain) is absent in *Chlorobaculum limnaeum* (freshwater). Nonetheless, both organisms contain the UDP-L-Qui2NAc pathway cluster.

### Prevalence of enantiomeric pairs and isomers of N-acetyl derivatives

Both enantiomers of a few monosaccharides are reported in natural glycans. The two enantiomers may or may not be biosynthesized from the same precursor, and may be linked to different nucleotides (Table S2). The present analysis shows that both enantiomers are found in only a small number of organisms, that too in specific genera, class or phyla (Table 3). Three isomeric N-acetyl derivatives of fucosamine (6-deoxygalactosamine) and of quinovosamine (6-deoxyglucosamine) are found in living systems. The N-acetyl group is present at C2, C3 or C4 position in these isomers. Only few organisms use more than one of these three isomers (Table 3). One such organism is *E. coli* NCTC11151 which contains both Fuc4NAc and Fuc3NAc. In contrast, *E. coli* O177:H21 uses L-Fuc2NAc along with Fuc3NAc. Genomic context analysis showed that Fuc4NAc biosynthesis genes are part of the O-antigen cluster in both these strains. On the other hand, genes for the biosynthesis of Fuc3NAc (in NCTC11151) and L-Fuc2NAc (in O177:H21) are present as part of the colanic acid cluster. Four genomes (strains) of *Pseudomonas orientalis* use Qui4NAc, Qui2NAc and L-Qui2NAc; genes required for the biosynthesis of these three monosaccharides are all in the same genomic neighbourhood.

**Table 3.** Presence of enantiomeric pairs and isomeric N-acetyl derivative pairs§.

| Monosaccharide | Where present | Number of organisms (genomes) in which these monosaccharide pairs are used |
| --- | --- | --- |
| **Both D- and L-enantiomers** |  |  |
| Galactose | Extremophiles | 33 |
| Fucose | Phylum FCB group | 7 |
|  | Phylum Deferribacteres | 1 |
|  | Class Gammaproteobacteria | 10 |
| Rhamnose | Genus *Pseudomonas* | 338 |
| 6-Deoxytalose | Genus *Pseudomonas* | 140 |
| Qui2NAc<sup>†</sup> | Phylum Proteobacteria | 64 |
| Fuc2NAc | Genus *Staphylococcus*<sup>¶</sup> | 300 |

| Isomers of N-acetyl derivatives |  |  |
| --- | --- | --- |
| Fuc3NAc and Fuc4NAc<sup>‡</sup> | Family Enterobacteriaceae | 33 |
| Qui2NAc, Qui4NAc | Several phyla | 193 |

<sup>§</sup> The diastereomeric pair of Fuc4NAc and L-Fuc2NAc are found in some strains of *Escherichia coli*.
<sup>¶</sup> Both Fuc2NAc and L-Fuc2NAc are components of capsular polysaccharides (27)
<sup>†</sup> Abbreviated names are used for some of the monosaccharides. Full names of these are given in Supplementary\_data.xlsx:Worksheet4.
<sup>‡</sup> Glucose-1-phosphate is the precursor for both Fuc3NAc and Fuc4NAc and UDP-Glc2NAc is the precursor for Fuc2NAc (the isomer that is absent in these organisms).

### Why are some pathways not found in Archaea?

Most of the rare group monosaccharides are absent in Archaea. Members of Euryarchaeota contain higher number of monosaccharides than TACK group. This could be suggestive of lateral gene transfer events with bacterial members as members of Euryarchaeota, particularly methanogens, coexist with other organisms in microbiomes (28) and have been inferred to acquire their genetic content (29). It is premature to associate absence of monosaccharide diversity to the apparent lack of pathogenicity in Archaea (28). This is because of inadequate information regarding the abundance of Archaea in various microbiomes. This in turn is due to our limitations in the detection of Archaea and associating them with disease phenotypes.

Apart from these possibilities, methodological limitations may have resulted in apparent absence of monosaccharides in Archaea. Only 4-5% of the 789 sequences used for generating HMM profiles or as BLASTp queries are from archaea. The pathway for the biosynthesis of TDP-/dTDP-L-rhamnose has four enzymes viz., RmlA, RmlB, RmlC and RmlD. Of these, only RmlB could not be found by HMM profile in *Saccharolobus sp., Desulfurococcus sp.* and *Sulfolobus sp*. leading to the conclusion that L-rhamnose is absent in these organisms. Analysis of the neighbourhood of RmlA, RmlC and RmlD revealed a sequence which could potentially be RmlB since it retains conserved residues of this family. This sequence could not be captured by the profile-based search due to stringent thresholds (=400 bits) [profile GPE05430; Supplementary_data.xlsx:Worksheet1]. Potential RmlB sequences of these organisms score 300-350 bits. This observation suggests that the pathway exists in these organisms but was not identified due to the stringency of the threshold. However, this is in contrast to other cases of absence of monosaccharides wherein none of the proteins of a pathway in the genome score even the default bit score of HMMER (i.e., 10 bits).

### Use of more than one nucleotide derivative/alternative pathways

L-rhamnose and Qui4NAc are biosynthesized as both UDP- and TDP-/dTDP-derivatives (Figures S2a, S2c). However, the TDP-/dTDP-pathways are found in Archaea and Bacteria, but not the UDP-pathways. TDP-/dTDP-6-deoxy-L-talose is biosynthesized via reduction of TDP-/dTDP-4-keto-L-rhamnose or C4 epimerization of TDP-/dTDP-L-rhamnose (Figure S2a). The former pathway occurs in 141 genomes belonging to multiple phyla and notably in *Pseudomonas sp., Streptococcus sp.* and *Streptomyces sp*. The latter pathway is found in 255 genomes belonging to Proteobacteria and Terrabacteria, and notably in *Burkholderia sp., Mycobacterium sp.* and *Xanthomonas oryzae*. N,N’-diacetyl legionaminic acid can be biosynthesized either from UDP-route or GDP-route (Figure S2f). The latter pathway is found in 93 of 96 genomes of *Campylobacter jejuni* whereas the former is found in 10 other genomes primarily belonging to Bacteriodetes/Chlorobi class.

## Discussion

The importance of glycans, especially in Archaea and Bacteria, is well documented. Establishing the specific role of glycans and studying structure-function relationship is largely hindered by factors such as non-availability of high-throughput sequencing methods, inadequate information as to which genes are involved in non-template driven biosynthesis, phase variation (30) and microheterogeneity (7). In this study, completely sequenced archaeal and bacterial genomes were searched for monosaccharide biosynthesis pathways using sequence homology-based approach. It is found that the usage of monosaccharides is not at all conserved across Archaea and Bacteria. This is in stark contrast to the alphabets of DNA and proteins which are universal. In addition, marked differences are observed even among different strains of a species. The range of monosaccharides used by an organism seems to be influenced by environmental factors such as growth (nutrients, pH, temperature, …) and environmental (host, microbiome, …) conditions. For instance, high uronic acid content in exopolysaccharides of marine bacteria imparts anionic property which is implicated in uptake of Fe^3+^ thus promoting its bioavailability to marine phytoplankton for primary production (31) and against degradation by microbes (32). Mutation in genes that encode enzymes for the biosynthesis of LPS in *E. coli* was shown to confer resistance to T7 phage (33). Thus, organisms, even at the level of strains, seem to evolve to modify their monosaccharide repertoire to increase fitness. In fact, selection pressure and horizontal gene transfer events could be the reason for the monosaccharide repertoire of bacteria far exceeding those of mammalian and other eukaryotes.

Genes encoding enzymes for the biosynthesis of Neu5Ac are found in 5% and 0.6% genomes of Alpha-proteobacteria and Actinobacteria, respectively; the bacterial carbohydrate structure database had no Neu5Ac-containing glycan from organisms belonging to this class/phylum (9). L-Rhamnose and L-fucose are found in 16% of Delta- and Epsilon-proteobacteria genomes and in 25% of actinobacteria genomes. However, very few L-rhamnose- and L-fucose-containing glycans from these classes/phyla are deposited in the database leading to the inference that these are rare sugars in this class/phylum. Thus, inferring monosaccharide usage based on an analysis of experimentally characterized glycans can at best give a partial picture.

Rare group monosaccharides are those which are found only in a few species, few genera and few phyla. Reasons for acquiring Rare group sugars can at best be speculative. For instance, Bac2Ac4Ac occurs at the reducing end of glycans N- and O-linked to proteins (34) but the presence of Bac2Ac4Ac is not mandatory for *C. jejuni* PglB, an oligosaccharyltransferase, since it can transfer glycans which have Glc2NAc, Gal2NAc or Fuc2NAc also at the reducing end (35). Perhaps, Bac2Ac4Ac provides resistance to enzymes like PNGase F that cleave off N-glycans. L-rhamnose, Neu5Ac, L-Qui2NAc, Man2NAc and L-Ara4N are not used by *Leptospira biflexa* (a non-pathogen) but are used by *Leptospira interrogans* (a pathogen). It is tempting to infer that these monosaccharides impart virulence to the latter but analysis of monosaccharides used by *E. coli* strains belonging to multiple pathotypes (enterohemorragic, enteropathogenic, uropathogenic) did not reveal any relationship between monosaccharides and their phenotype. Tyvelose, paratose and abequose are 3,6-dideoxy sugars that belong to the Rare group. These are found primarily in *Salmonella enterica, Yersinia pestis* and *Yersinia pseudotuberculosis*. These are present in the O-antigen of *Y. pseudotuberculosis* (36). *Y. pestis*, closely related to and derived from *Y. pseudotuberculosis*, lacks O-antigen (rough phenotype) due to the silencing of O-antigen cluster (37). *Y. enterocolitica*, also an enteric pathogen like *Y. pseudotuberculosis*, does not contain these monosaccharides. Hence the role of these 3,6-dideoxy sugars in the O-antigen of *Y. pseudotuberculosis* does not seem to be related to enteropathogenicity.

Besides answering the question of the universality of glycan alphabet, this study also has led to certain beneficial outcomes. L-rhamnose, mannose and L-Pse5ac7Ac are found in *B. cereus, B. mycoides* and *B. thuringeinsis* but not in *B. subtilis, B. amyloliquefaciens, B. licheniformis, B. velezensis and B. vallismortis.* Such differences may be potentially be exploited towards taxonomic identification, provided that these patterns hold true after analysis of a larger number of strains from each of these species. Enzymes synthesizing monosaccharides that are exclusive to a pathogen vis-à-vis its host can be identified as potential drug targets. An illustrative example is of the non-hydrolyzing C2 epimerase: it mediates the synthesis of UDP-Man2NAc, UDP-L-Qui2NAc, UDP-L-Fuc2NAc and UDP-Man2NAc3NAc and is found in 60% of the archaeal+bacterial genomes but not in humans (human genome was scanned for the presence of these pathways; unpublished results). It has already been reported that inhibitors of this enzyme are effective against methicillin-resistant *S. aureus* and a few other bacteria (38). Based on the prevalence of this enzyme in all other phyla, inhibitors against this enzyme would be promising broad spectrum antimicrobial therapies. As already noted (39), knowledge of monosaccharide composition is also useful in ensuring consistency of recombinant glycoprotein therapeutics. Knowledge of biosynthesis pathways also allows cloning the entire cassette in a heterologous host for large scale production of monosaccharides for commercial and research applications.

Thus, glycans show least evolutionary conservation among these three macromolecules (40). Owing to their virtue of endowing distinction, existence of a universal glycan alphabet is antithetical. Here, alphabet is used in the same sense as its dictionary meaning, viz., a set of letters or symbols which combine to form complex entities. In the case of glycans, structural diversity arises not only by the set of monosaccharides an organism uses but also by linkage variations (α1→3, β1→4, etc.), branching and modifications (e.g., sulfation, acetylation, …). Knowledge of the linkage types, branching patterns and modifications that an organism uses will further our understanding of the biological roles of glycans.

## Author contributions

PVB conceived and supervised the research, PS conceived the design and guided the development of GlycoPathDB. JS performed the research and developed the database. PVB and JS wrote the paper.

## Conflicts of interest

The author(s) declare that there are no conflicts of interest

## Funding information

This work received no specific grant from any funding agency

## Acknowledgments

We thank Nitesh Kumar, Ruchi Kumari, Tejas Shah and Tejas Vaidya for technical assistance on the development of GlycoPathDB. We thank Shradha Khater and Toshi Mishra for useful discussion. Jaya Srivastava is thankful to the Council of Scientific and Industrial Research, Government of India for research fellowship (File number 09/087/(0877)/2017-EMR-I).

## Notes

http://www.bio.iitb.ac.in/glycopathdb/

### Competing Interest Statement

The authors have declared no competing interest.

### Summary of Updates

Changes introduced in analysis of monosaccharides in Archaea in results section. Minor revision in usage of terms and conclusions also included

