## supplementary material for "The glycan alphabet is not universal: a hypothesis"

### CONTENTS


| Data | Description |
|---|---|
| Figure S1 | Number of organisms with different number of strains sequenced |
| Figure S2 | Biosynthesis pathways |
| Figure S3 | Proteome sizes for different number of monosaccharides |
| Figure S4 | Prevalence of monosaccharides in species versus that in genomes |
| Figure S5 | Bit score distribution plots for hits of various pairs of profiles |
| Table S1 | Tools and databases used in this study |
| References | References cited in Table S1 |
| Table S2 | Comparison of the precursor and nucleotide used for the biosynthesis of two enantiomers of a monosaccharide |
| Flowchart S1 | Procedure used to generate HMM profiles |
| Flowchart S2 | Precedence rules for assigning annotation to proteins that are hits to two or more profiles and/or BLASTp queries |
| References | References to the research articles which describe the pathways (or enzymes of the pathways) of monosaccharide biosynthesis. These formed the basis for generating HMM profiles and choosing BLASTp queries. |


**MS-EXCEL file provided separately:** Supplementary Data.xlsx


| | |
|---|---|
| Worksheet1 | Details of HMM profiles |
| Worksheet2 | Details of BLASTp queries |
| Worksheet3 | Prevalence of monosaccharides in genomes / species |
| Worksheet4 | Abbreviated names of monosaccharides |
| Worksheet5 | Enzyme types, enzymes and monosaccharide groups |
| Worksheet6 | Precursors of various monosaccharides |

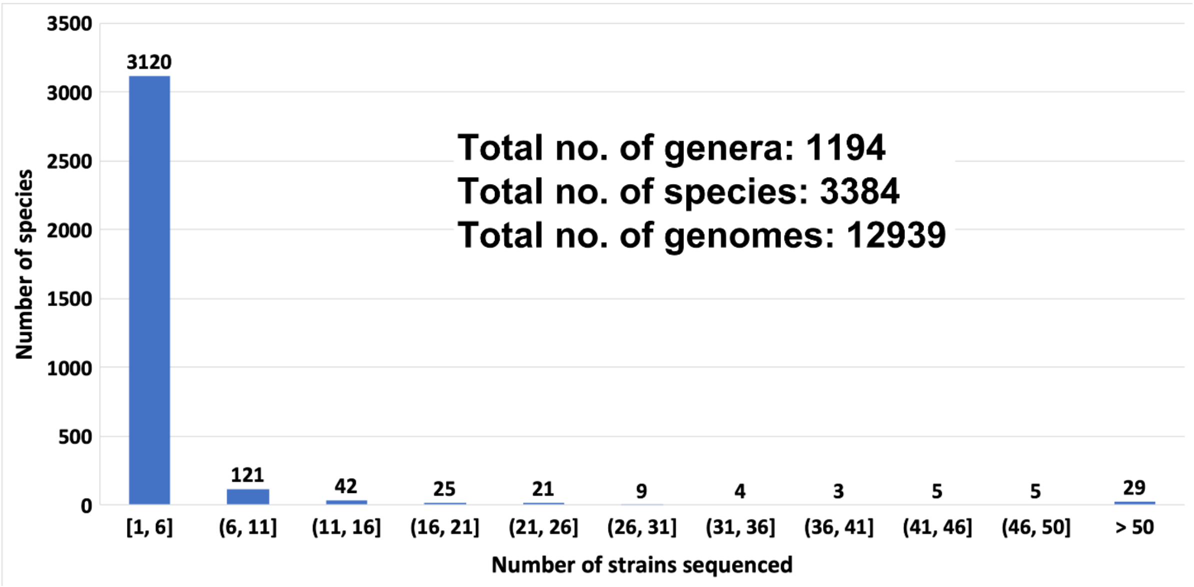


**Figure S1** The number of species for which different number of strains are sequenced. Six or fewer strains are sequenced for most of the species. On the other hand, more than 50 strains are sequenced for 29 species. *Escherichia coli* and *Salmonella enterica* have the highest number of sequenced strains (714 and 602, respectively). Genus and species names are not known for 45 endosymbionts; only their host name is known e.g., Legionella endosymbiont. Each such case is considered as a distinct species.

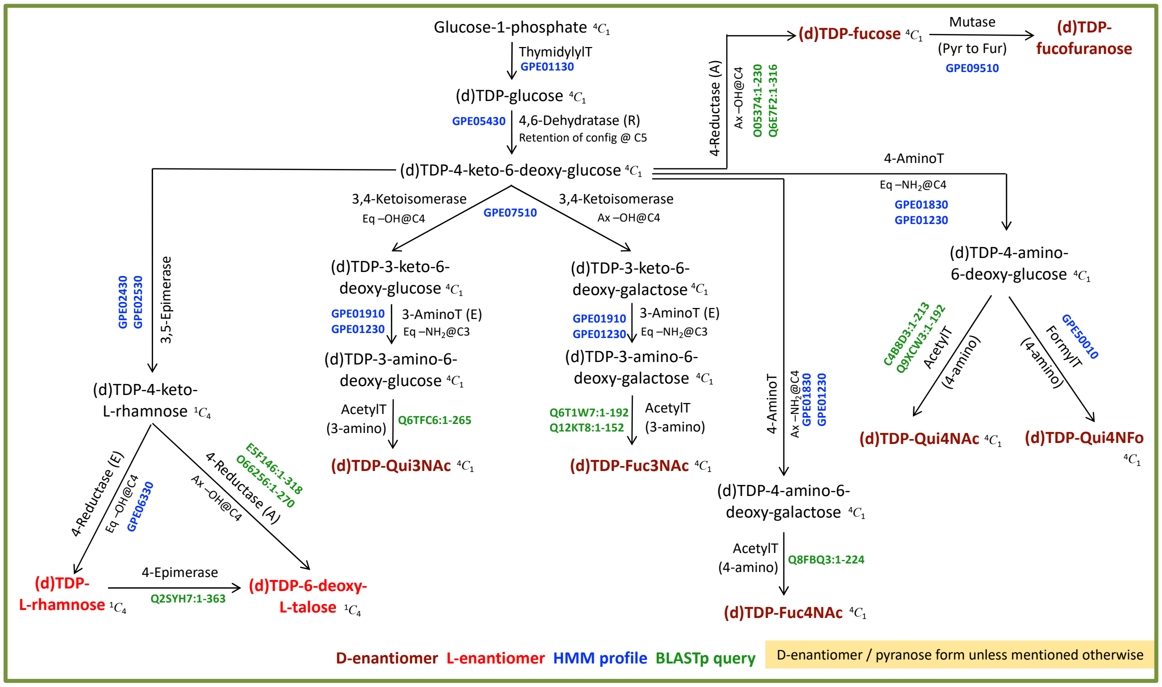


**Figure S2a** TDP-/dTDP-linked monosaccharides derived from glucose-1-phosphate. Abbreviated names are used for some of the monosaccharides. Full names of these are given in Supplementary_data.xlsx:Worksheet4.

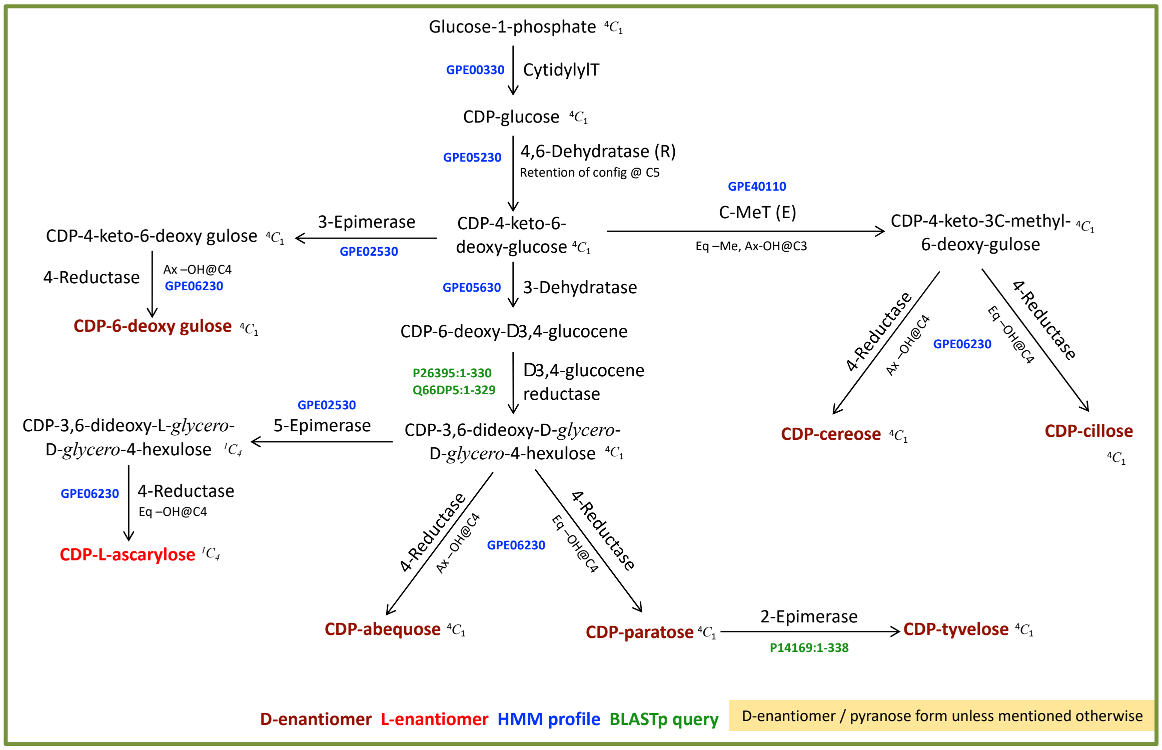


**Figure S2b** CDP-linked monosaccharides derived from glucose-1-phosphate. Abbreviated names are used for some of the monosaccharides. Full names of these are given in Supplementary_data.xlsx:Worksheet4.

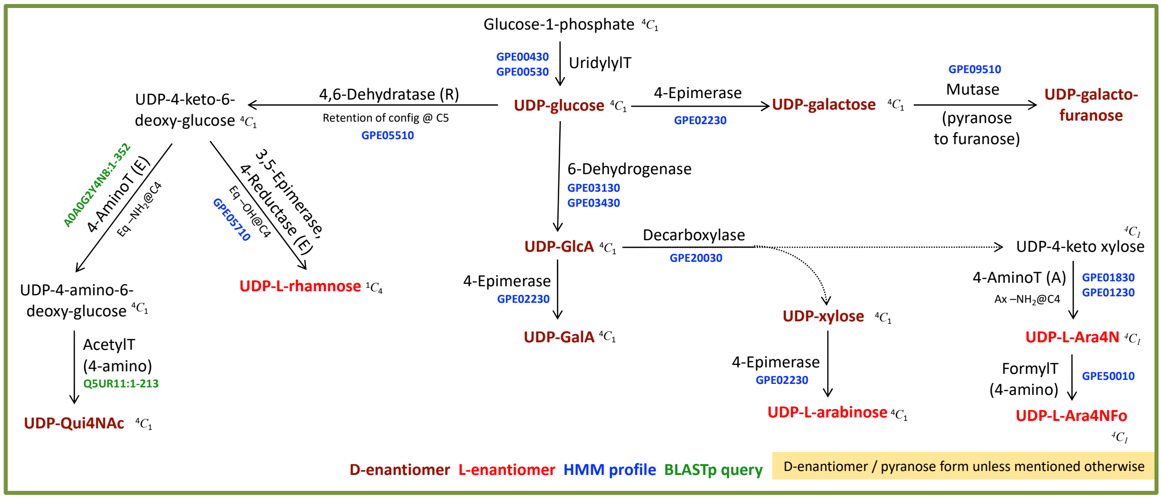


**Figure S2c** UDP-linked monosaccharides derived from glucose-1-phosphate. Abbreviated names are used for some of the monosaccharides. Full names of these are given in Supplementary_data.xlsx:Worksheet4.

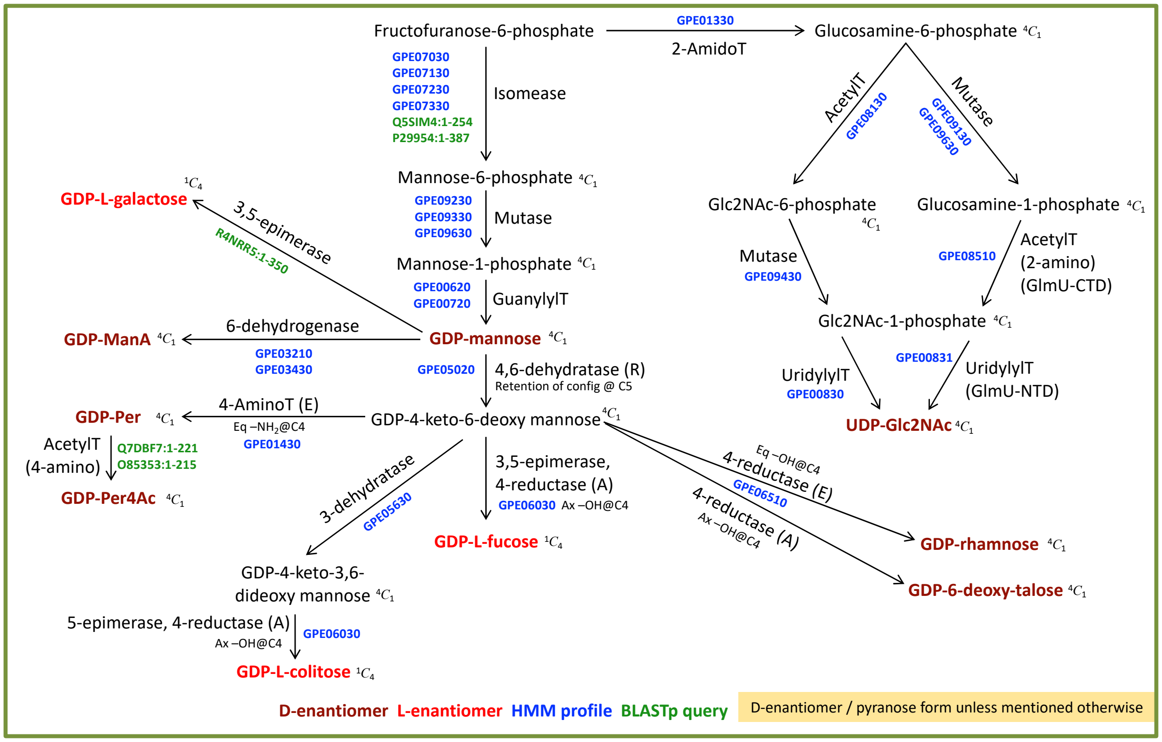


**Figure S2d** GDP- and UDP-linked monosaccharides derived from fructofuranose-6-phosphate. Abbreviated names are used for some of the monosaccharides. Full names of these are given in Supplementary_data.xlsx:Worksheet4.

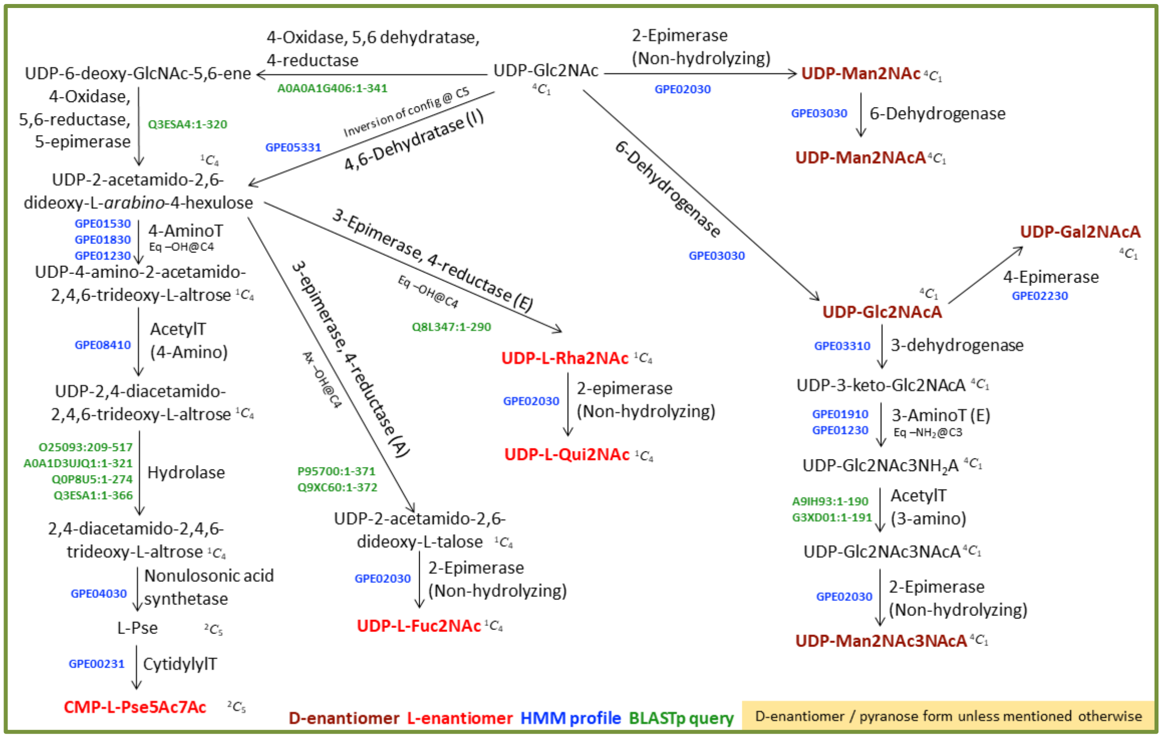


**Figure S2e** CMP- and UDP-linked monosaccharides derived from UDP-Glc2NAc (1 of 2). Abbreviated names are used for some of the monosaccharides. Full names of these are given in Supplementary_data.xlsx:Worksheet4.

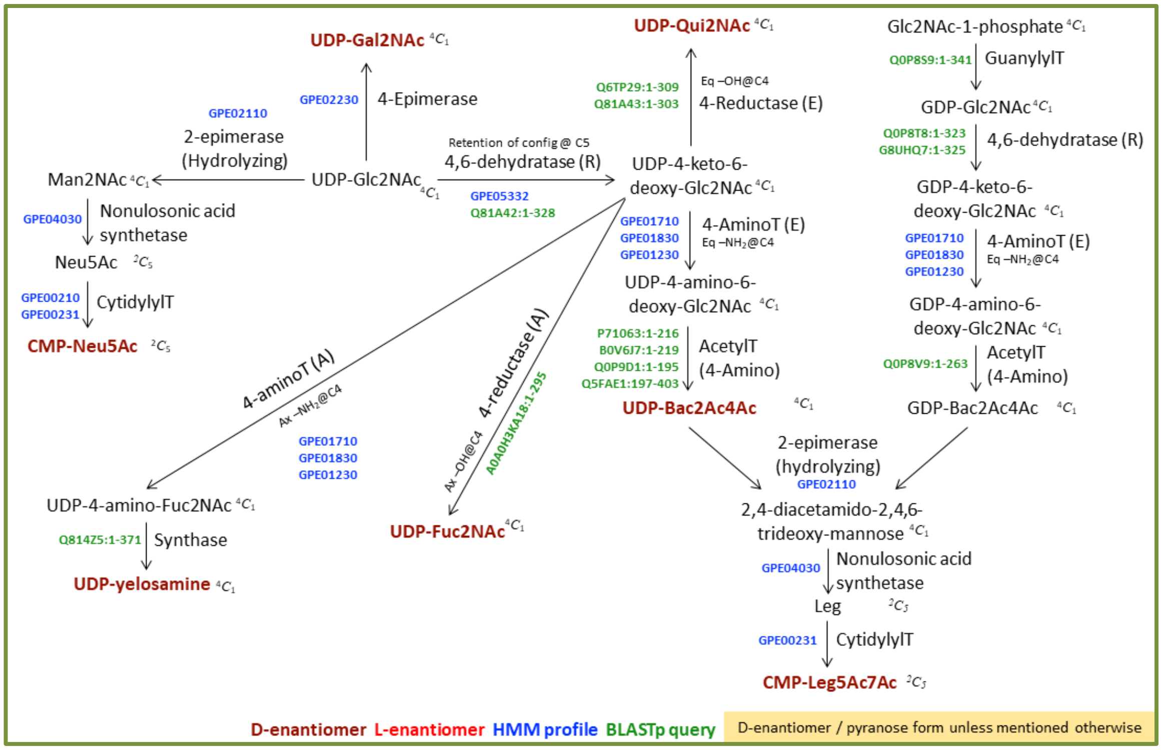


**Figure S2f** CMP- and UDP-linked monosaccharides derived from UDP-Glc2NAc (2 of 2). CMP-Leg5Ac7Ac may be biosynthesized through GDP-linked or UDP-linked intermediates. Abbreviated names are used for some of the monosaccharides. Full names of these are given in Supplementary_data.xlsx:Worksheet4.

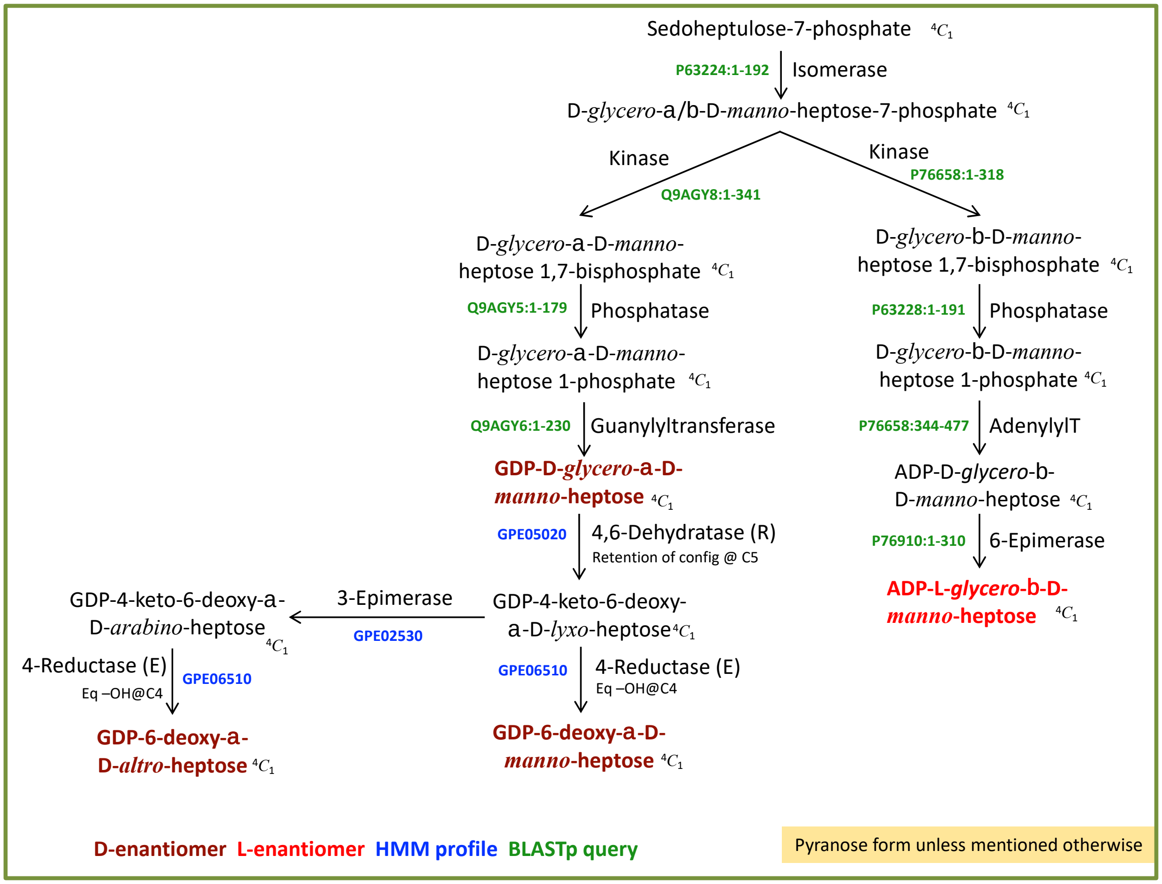


**Figure S2g** ADP- and GDP-linked heptoses derived from sedoheptulose-7-phosphate. Abbreviated names are used for some of the monosaccharides. Full names of these are given in Supplementary_data.xlsx:Worksheet4.

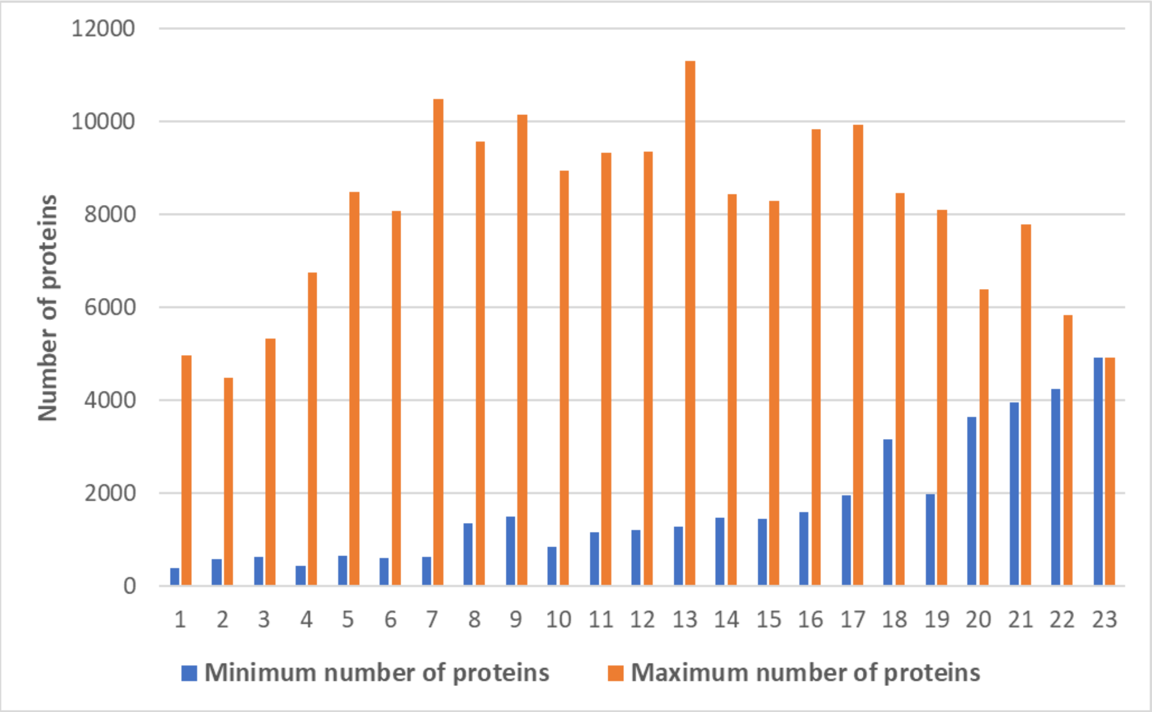


**Figure S3** Variations in the proteome size of organisms which encode the same number of monosaccharides. Only the smallest and largest proteome sizes are shown. As can be seen, the number of monosaccharides used by an organism is independent of the proteome size. For instance, *Helicobacter pylori* PNG84A (proteome size = 1353) uses the same number of monosaccharides (7) as *Sorangium cellulosum* So0157-2 (proteome size = 10480).

62

**A**

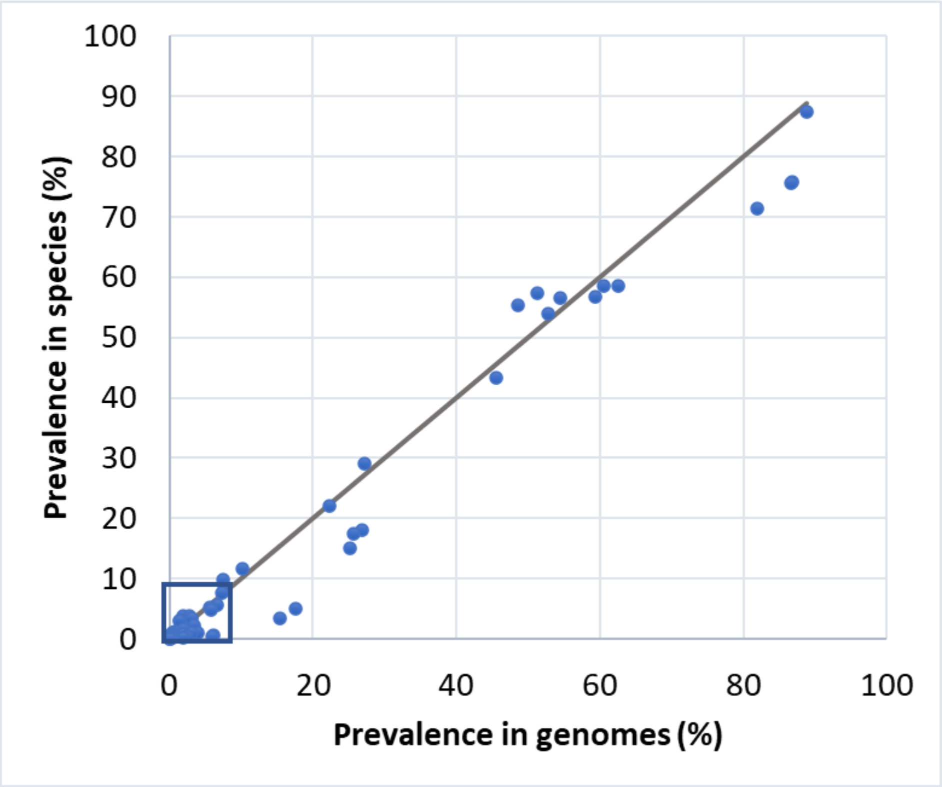

Prevalence in species (%)
Prevalence in genomes (%)
0
20
40
60
80
100
10
30
50
70
90


**B**

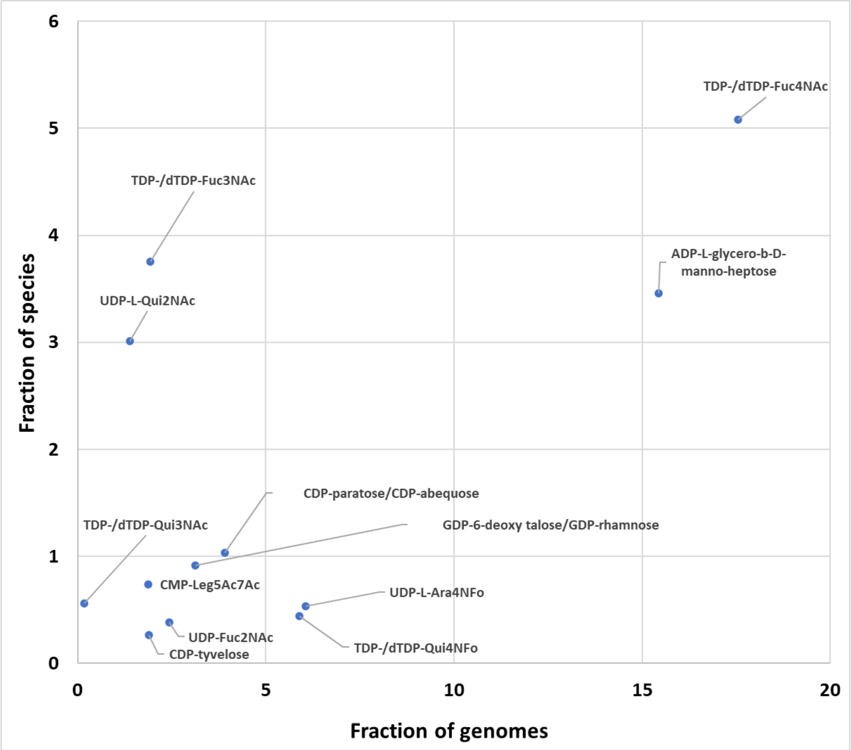

Fraction of species
Fraction of genomes
TDP-/dTDP-Fuc4NAc
TDP-/dTDP-Fuc3NAc
ADP-L-glycero-b-D-manno-heptose
UDP-L-Qui2NAc
CDP-paratose/CDP-abequose
GDP-6-deoxy talose/GDP-rhamnose
TDP-/dTDP-Qui3NAc
CMP-Leg5Ac7Ac
UDP-L-Ara4NFo
UDP-Fuc2NAc
CDP-tyvelose
TDP-/dTDP-Qui4NFo

**Figure S4** (A) The prevalence of each monosaccharide as percentages of the genomes analyzed in this study (viz., 12939) and the number of species covered by these genomes (viz., 3384; Figure S1(a)). The diagonal line is manually drawn to facilitate visualization of deviations. (B) Zoomed in view of the region near the origin in (A). Data for most of the monosaccharides lie on the diagonal suggesting that the sequencing of a large number of strains for a few species has not biased the outcome, with the exception of TDP-/dTDP-Fuc4NAc and UDP-L-Qui2NAc. TDP-/dTDP-Fuc4NAc (a point below the diagonal line) is present in fewer species but represents a larger fraction of genomes since 679 strains of *E. coli* contain this monosaccharide. Conversely, presence of UDP-L-Qui2NAc (a point above the diagonal line) is highly strain specific. Abbreviated names are used for some of the monosaccharides. Full names of these are given in Supplementary_data.xlsx:Worksheet4.

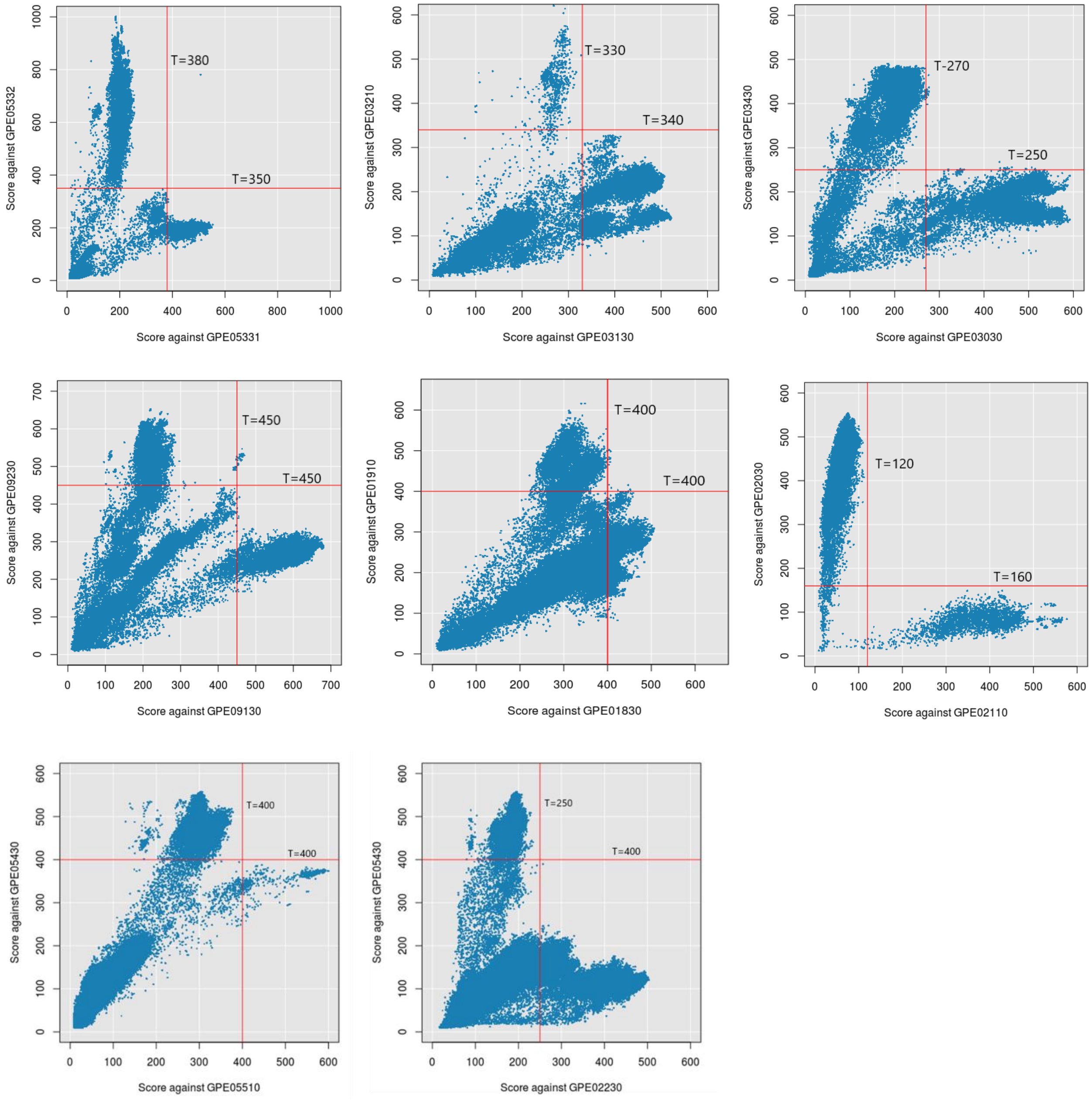


**Figure S5** Setting bit score thresholds for HMM profiles with varying substrate specificities. TrEMBL database was scanned using the profiles shown along the X- and Y-axes in the above scatter plots; for these scans, default values set by HMMer were used for all the parameters. Hits that are common to a pair of profiles (shown along X- and Y-axes) were chosen and bit scores of such hits were plotted against each other. Bit score thresholds (indicated by red lines) were chosen such that a protein is a hit for only one of the two profiles. Threshold was revised for GPE05331 set to exclude PdeG.

**Table S1 Tools and databases used in this study**

| Tool / Database | Version / Release | URL | Reference |
|---|---|---|---|
| Tools installed and run locally on a Linux platform | | | |
| BLASTp | 2.2.31+ | ftp://ftp.ncbi.nlm.nih.gov/blast/executables/blast+/2.2.31/ | (1) |
| HMMER | 3.1b2 | http://hmmer.org/download.html | (2) |
| MUSCLE | 3.8.31 | https://www.ebi.ac.uk/Tools/msa/muscle/ | (3) |
| CD-Hit | 4.6 | http://weizhongli-lab.org/cd-hit/ | (4) |
| Directly accessed from the website or FTP site | | | |
| UniProt | 2018_07 | https://www.uniprot.org/ | (5) |
| Genome | 2019_03 | https://www.ncbi.nlm.nih.gov/genome/ | (6) |
| Pubmed | Not applicable | https://www.ncbi.nlm.nih.gov/pubmed/ | (6) |
| CATH-Plus | 4.2 | http://www.cathdb.info/ | (7) |
| PDB | Not applicable | https://www.rcsb.org/ | (8) |
| Used through the TrEMBL database | | | |
| UniRule | Not applicable | https://www.uniprot.org/help/unirule | (5) |
| SAAS | Not applicable | https://www.uniprot.org/help/saas | (5) |

**Table S2** Monosaccharides whose both enantiomers are considered in the present study: comparison of the nucleotide to which the enantiomer is linked and the precursor for its biosynthesis

| Monosaccharide | D enantiomer | | L enantiomer | |
|---|---|---|---|---|
| | Nucleotide | Precursor | Nucleotide | Precursor |
| Rhamnose | GDP | Glc-1-P | TDP, dTDP, UDP | Glc-1-P |
| 6-Deoxytalose | GDP | Glc-1-P | TDP, dTDP | Glc-1-P |
| Galactose | UDP | Glc-1-P | GDP | Glc-1-P |
| Fucose | TDP, dTDP | Glc-1-P | GDP | Glc-1-P |
| Fuc2NAc | UDP | UDP-Glc2NAc | UDP | UDP-Glc2NAc |
| Qui2NAc | UDP | UDP-Glc2NAc | UDP | UDP-Glc2NAc |

**Flowchart S1** Procedure used to generate HMM profiles

| | |
|---|---|
| 1. Generation of Exp dataset and Exp profile, and setting $T_{exp}$ | |
| Step 1a | Consider only those enzymes which are characterized by direct enzyme activity assay |
| Step 1b | Remove redundancy (80% sequence identity cutoff) and obtain a multiple sequence alignment (MSA) |
| Step 1c | Use the MSA as input to generate an HMM profile |
| Step 1d | Score Exp dataset sequences against this HMM profile |
| Step 1e | Set the bit score of the lowest scoring sequence as the bit score threshold for Exp dataset, $T_{exp}$ |
| 2. Generation of Extend dataset and Extend profile, and setting $T_{extend}$ | |
| Step 2a | Add sequences that meet any of the following criteria to the Exp dataset |
| (i) | SwissProt entries satisfying the threshold $T_{exp}$ |
| (ii) | SwissProt entries scoring $< T_{exp}$ provided they show conservation of active site residues. Active site residues were collated based on site directed mutagenesis studies or ligand-bound 3D structures |
| (iii) | TrEMBL entries for which molecular function has been inferred from experiments other than direct enzyme assays viz., complementation assays, phenotypic studies, etc. |
| (iv) | TrEMBL entries with solved 3D structure |
| (v) | FunFam members (CATH database) but only in the case of CDP-glucose 4,6-dehydratase (FunFam 20603) and phosphomannoisomerase family 3 (FunFam 54112) |
| Step 2b | Remove redundancy (80% sequence identity cutoff) and obtain a multiple sequence alignment (MSA) |
| Step 2c | Use the MSA as input to generate an HMM profile |
| Step 2d | Score Extend dataset sequences against this HMM profile |
| Step 2e | Set the bit score of the lowest scoring sequence as the bit score threshold for Extend dataset, $T_{extend}$ |

**Flowchart S2** Precedence rules for assigning annotation to proteins that are hits to two or more profiles and/or BLASTp queries

```
# Case 1 of 14
# specific_aminoTs = [GPE01710, GPE01430, GPE01530]
# C3_C4_aminoTs = [GPE01910, GPE01830]
# EXPECTED: for a protein which is a hit for one of the specific_aminoTs is expected
   to be a hit in C3_C4_aminoTs as well as GPE01230

    IF (hit for any one of specific_aminoTs) THEN
      IF (hit for any one of C3_C4_aminoTs) THEN
         IF (hit for GPE01230) THEN
            Pass (i.e., this is as expected)
         ELSE
            Alert: Hit for one of C3_C4_aminoTs but not GPE01230
         ENDIF
      ELSE
         Alert: Hit for one of specific_aminoTs but not C3_C4_aminoTs
      ENDIF
    ENDIF
    IF (hit for any one of C3_C4_aminoTs) THEN
      IF (hit for GPE01230) THEN
         Pass (as expected)
      ELSE
         Alert: Hit for one of C3_C4_aminoTs but not GPE01230
      ENDIF
   ENDIF
```

```
# Case 2 of 14
   IF a protein is a hit for any one of specific_aminoTs, it should be assigned that
   annotation
       IF (hit for any one of specific_aminoTs) THEN
         Assign annotation
       ENDIF
   ENDIF
```

```
# Case 3 of 14
# For a protein which is a hit for one of C3_C4_aminoTs but not any of
specific_aminoTs, it should be assigned the former
    IF (hit for one of C3_C4_aminoTs and not for any of specific_aminoTs) THEN
      Assign GPE01830/GPE01910 annotation
    ENDIF
```

```
# Case 4 of 14
# For a protein which is a hit for GPE00210
# GPE00210 is used only in combination with GPE00231
# A protein which is a hit for GPE00210 is expected to be a hit for GPE00231 also
    IF (hit for GPE00210) THEN
      IF (hit for GPE00231) THEN
         Assign GPE00210 annotation
      ELSE
         Alert: Hit for GPE00210 but not GPE00231
      ENDIF
    ENDIF
```

```
# Case 5 of 14
# For a protein which is a hit for GPE02430
# GPE02430 is used only in combination with GPE02530
# A protein which is a hit for GPE02430 is expected to be a hit for GPE02530 also
    IF (hit for GPE02430) THEN
      IF (hit for GPE002530) THEN
         Assign GPE02430 annotation
      ELSE
         Alert: Hit for GPE02430 but not GPE02530
      ENDIF
    ENDIF
```

```
# Case 6 of 14
# For a protein that is a hit for GPE03130
# GPE03130 is used only in combination with GPE03430
# A protein which is a hit for GPE03130 is expected to be a hit for GPE03430 also
    IF (hit for GPE03130) THEN
      IF (hit for GPE03430) THEN
         Assign GPE03130 annotation
      ELSE
         Alert: Hit for GPE03130 but not GPE03430
      ENDIF
    ENDIF
```

```
# Case 7 of 14
# For a protein that is a hit for GPE03210
# GPE03210 is used only in combination with GPE03430
# A protein which is a hit for GPE03210 is expected to be a hit for GPE03430 also
    IF (hit for GPE03210) THEN
      IF (hit for GPE03430) THEN
          Assign GPE03210 annotation
      ELSE
          Alert: Hit for GPE03210 but not GPE03430
      ENDIF
    ENDIF
```

```
# Case 8 of 14
# For a protein that is a hit for GPE09130
# GPE09130 is used only in combination with GPE09630
# A protein which is a hit for GPE09130 is expected to be a hit for GPE09630 also
    IF (hit for GPE09130) THEN
      IF (hit for GPE09630) THEN
          Assign GPE09130 annotation
      ELSE
          Alert: Hit for GPE09130 but not GPE09630
      ENDIF
    ENDIF
```

```
# Case 9 of 14
# For a protein that is a hit for GPE09230
# GPE09230 is used only in combination with GPE09330 and GPE09630
# A protein which is a hit for GPE09230 is expected to be a hit for GPE09630 also
    IF (hit for GPE09230) THEN
      IF (hit for GPE09630) THEN
          Assign GPE09230 annotation
      ELSE
          Alert: Hit for GPE09230 but not GPE09630
      ENDIF
    ENDIF
```

```
# Case 10 of 14
# For a protein that is a hit for GPE09330 and GPE09630
# GPE09330 is used only in combination with GPE09630
# A protein can be a hit for GPE09330 or GPE09630, but not for both (non-
  orthologous)
    IF (hit for GPE09330 AND hit for GPE09630) THEN
        Alert: Hit for GPE09330 and GPE09630
    ENDIF
```

```
# Case 11 of 14
# For a protein that is a hit for GPE00620 and GPE00720
# GPE00620 is used only in combination with GPE00720
# A protein can be a hit for GPE00620 or GPE00720, but not for both (non-
  orthologous)
    IF (hit for GPE00620 AND hit for GPE00720) THEN
        Alert: Hit for GPE00620 and GPE00720
    ENDIF
```

```
# Case 12 of 14
# Isomerases: GPE07030, GPE07130, GPE07230, and GPE07330
# A protein can be a hit for any one of the above four profiles (non-orthologous)
      For GPE07030, GPE07130, GPE07230 and GPE07330
      IF (hit for more than one)
           Alert: Hit for (list all profiles which appear as hits from above list)]
```

```
# Case 13 of 14
# For a protein that is a hit for GPE00430 and GPE00530
# GPE00430 is used only in combination with GPE00530
# A protein can be a hit for GPE00430 or GPE00530, but not for both (non-
  orthologous)
    IF (hit for GPE00430 AND hit for GPE00530) THEN
        Alert: Hit for GPE00430 and GPE00530
    ENDIF
```

```
# Case 14 of 14
# For a protein that is a hit for GPE05332 and Q81A42:1-328
# GPE05332 is used only in combination with Q81A42:1-328
# A protein can be a hit for GPE05332 or Q81A42:1-328, but not for both (non-
  orthologous)
    IF (hit for GPE05332 AND hit for Q81A42:1-328) THEN
        Alert: Hit for GPE05332 and Q81A42:1-328
    ENDIF
```

114

Research articles which report the characterization of enzymes involved in the biosynthesis of monosaccharides are listed below. Amino acid sequences of these enzymes were either used to generate HMM profiles or used as BLASTp queries. The PubMed Ids of these research articles are included in the GlycoPathDB (www.bio.iitb.ac.in/glycopathdb/) against respective sequence entry. These PubMed Ids are hyperlinked to the corresponding PubMed webpage.

10. Jia X, Kang J, Yin H. A simple and rapid method for measuring α-D-phosphohexomutases activity by using  anion-exchange chromatography coupled

with an electrochemical detector. PeerJ. 2016;4:e1517.

32. Thoden JB, Holden HM. Biochemical and structural characterization of WlbA

from Bordetella pertussis and Chromobacterium violaceum: enzymes required for the biosynthesis of 2,3-diacetamido-2,3-dideoxy-D-mannuronic acid. Biochemistry. 2011 Mar 8;50(9):1483–91.

54. Gu X, Lee SG, Bar-Peled M. Biosynthesis of UDP-xylose and UDP-arabinose in Sinorhizobium meliloti 1021: first characterization of a bacterial UDP-xylose

synthase, and UDP-xylose 4-epimerase. Microbiology. 2011 Jan;157(Pt 1):260–9.

65. Tenhaken R, Voglas E, Cock JM, Neu V, Huber CG. Characterization of GDP-mannose dehydrogenase from the brown alga Ectocarpus siliculosus providing the

precursor for the alginate polymer. J Biol Chem. 2011 May 13;286(19):16707–15.

88. Roper JR, Ferguson MAJ. Cloning and characterisation of the UDP-glucose 4’-

epimerase of Trypanosoma cruzi. Mol Biochem Parasitol. 2003 Nov;132(1):47–53.

99. Suryanti V, Nelson A, Berry A. Cloning, over-expression, purification, and

characterisation of N-acetylneuraminate synthase from Streptococcus agalactiae. Protein Expr Purif. 2003 Feb;27(2):346–56.

110. Riegler H, Herter T, Grishkovskaya I, Lude A, Ryngajllo M, Bolger ME, et al. Crystal structure and functional characterization of a glucosamine-6-phosphate.

Biochem J. 2012 Apr 15;443(2):427–37.

175. Piacente F, De Castro C, Jeudy S, Molinaro A, Salis A, Damonte G, et al. Giant virus Megavirus chilensis encodes the biosynthetic pathway for uncommon

acetamido sugars. J Biol Chem. 2014 Aug 29;289(35):24428–39.

209. Park NY, Lee JH, Kim MW, Jeong HG, Lee BC, Kim TS, et al. Identification of the Vibrio vulnificus wbpP gene and evaluation of its role in virulence. Infect

Immun. 2006 Jan;74(1):721–8.

218. Chen H, Zhao Z, Hallis TM, Guo Z, Liu Hw H. Insights into the Branched-Chain Formation of Mycarose: Methylation Catalyzed by an (S)-Adenosylmethionine-Dependent Methyltransferase We are grateful to Dr. Eugene Seno and the Lilly Research Laboratories for their generous gift of the plasmid pHJL311 and to the National Institutes of Health for grants (GM 35 906 and 54 346). H.-w.L. also thanks the National Institute of General Medical Sciences for a MERIT Award. T.M.H. was a trainee of the National Institute of General Medical Sciences (Biotechnology Training Grant: 2 T32 GM08347). Angew Chem Int Ed Engl. 2001 Feb 2;40(3):607–10.

219. Hofmeister DL, Thoden JB, Holden HM. Investigation of a sugar N-formyltransferase from the plant pathogen Pantoea ananatis. Protein Sci. 2019

Apr;28(4):707–16.

253. Babaoglu K, Page MA, Jones VC, McNeil MR, Dong C, Naismith JH, et al. Novel inhibitors of an emerging target in Mycobacterium tuberculosis; substituted

thiazolidinones as inhibitors of dTDP-rhamnose synthesis. Bioorg Med Chem Lett. 2003 Oct 6;13(19):3227–30.

274. Bruender NA, Holden HM. Probing the catalytic mechanism of a C-3’-methyltransferase involved in the biosynthesis of D-tetronitrose. Protein Sci. 2012

Jun;21(6):876–86.

285. Ding L, Seto BL, Ahmed SA, Coleman WGJ. Purification and properties of the Escherichia coli K-12 NAD-dependent nucleotide diphosphosugar epimerase,

ADP-L-glycero-D-mannoheptose 6-epimerase. J Biol Chem. 1994 Sep 30;269(39):24384–90.

296. Giraud MF, Leonard GA, Field RA, Berlind C, Naismith JH. RmlC, the third enzyme of dTDP-L-rhamnose pathway, is a new class of epimerase. Nat Struct

Biol. 2000 May;7(5):398–402.

308. Chantigian DP, Thoden JB, Holden HM. Structural and biochemical

characterization of a bifunctional ketoisomerase/N-acetyltransferase from Shewanella denitrificans. Biochemistry. 2013 Nov 19;52(46):8374–85.

318. Gunawan J, Simard D, Gilbert M, Lovering AL, Wakarchuk WW, Tanner ME, et al. Structural and mechanistic analysis of sialic acid synthase NeuB from Neisseria

meningitidis in complex with Mn2+, phosphoenolpyruvate, and N-acetylmannosaminitol. J Biol Chem. 2005 Feb 4;280(5):3555–63.

363. Vann WF, Daines DA, Murkin AS, Tanner ME, Chaffin DO, Rubens CE, et al.

The NeuC protein of Escherichia coli K1 is a UDP N-acetylglucosamine 2-epimerase. J Bacteriol. 2004 Feb;186(3):706–12.

374. Kepes F, Schekman R. The yeast SEC53 gene encodes phosphomannomutase. J

Biol Chem. 1988 Jul 5;263(19):9155–61.

385. Rösti J, Barton CJ, Albrecht S, Dupree P, Pauly M, Findlay K, et al. UDP-glucose 4-epimerase isoforms UGE2 and UGE4 cooperate in providing UDP-galactose for

cell wall biosynthesis and growth of Arabidopsis thaliana. Plant Cell. 2007 May;19(5):1565–79.
